# Expanding heterochromatin reveals discrete subtelomeric domains delimited by chromatin landscape transitions

**DOI:** 10.1101/187138

**Authors:** Antoine Hocher, Myriam Ruault, Petra Kaferle, Marc Descrimes, Mickael Garnier, Antonin Morillon, Angela Taddei

## Abstract

The eukaryotic genome is divided into chromosomal domains of heterochromatin and euchromatin. Transcriptionally silent heterochromatin is found at subtelomeric regions, leading to the telomeric position effect (TPE) in yeast, fly and man. Heterochromatin generally initiates and spreads from defined loci, and diverse mechanisms prevent the ectopic spread of heterochromatin into euchromatin. Here, we overexpressed the silencing factor Sir3 at various levels in yeast, and found that Sir3 spreading into Extended Silent Domains (ESD) eventually reached saturation at subtelomeres. We observed that Sir3 spreading into ESDs covered zone associated with specific histone marks in wild-type cells and stopped at zones of histone mark transitions including H3K79 tri-methylation levels. The conserved enzyme Dot1 deposits H3K79 methylation, and we found that it is essential for viability upon overexpression of Sir3, but not of a spreading-defective mutant Sir3A2Q. These data suggest that H3K79 methylation actively blocks Sir3 spreading. Lastly, our meta-analysis uncovers previously uncharacterized discrete subtelomeric domains associated with specific chromatin features offering a new viewpoint on how to separate subtelomeres from the core chromosome.

## Introduction

Heterochromatin classically designates chromosomal domains that remain condensed throughout the cell cycle (1). It impacts many aspects of chromosome biology, including genomic stability and gene expression (2). In contrast to gene specific repressors, heterochromatin silences genes independently of their DNA sequence (3). Heterochromatin is prevalent in eukaryotic genomes and is a major system of gene regulation, key to processes ranging from gene dosage compensation to cell differentiation to speciation (2).

In many species heterochromatin is associated to telomeres and to a portion of subtelomeres(4). Subtelomeres are genomic domains particularly difficult to define. While they often exhibit structural and functional properties, such as the presence of specific gene families, chromatin marks or a relatively fast gene turnover, there is no strict definition that segregates all these properties between subtelomeres and the core genome (4).

Transcriptional silencing generally initiates at defined loci and propagates by self-recruitment mechanisms (2,5–7). The coupling of histone modifying enzymes to nucleosomes allows the specific binding of silencing effectors and drives the formation of heterochromatin domains (8,9). However, the spread of heterochromatin must be limited to prevent encroaching on euchromatin (10).

In budding yeast, the silent information regulator (SIR) proteins, Sir2 Sir3 and Sir4, implement stable repression of cryptic mating type loci (HM) and semi-stable repression of genes near telomeres (7,11–16). The SIR complex is recruited at these loci by combination of specific DNA binding proteins that have functions outside silencing. At telomeres, the Repressor activator Rap1 binds the degenerated telomeric sequence TG1–3 (17), and recruits the SIR complex through direct interaction with Sir3 and Sir4. This recruitment is reinforced by additional interactions between Sir4 and the Ku heterodimers (18,19) Once nucleated, the activity of Sir2, a conserved histone deacetylase, creates favorable binding sites for Sir3, which preferentially binds deacetylated H4K16. Iterative cycles of Sir2-mediated histone deacetylation and Sir3-binding allow the self-propagation of the SIR complex on chromatin until a barrier is eventually reached (7,11).

Boundaries restrict silent domains at the mating type loci (20,21). A tRNA confines the Sir complex to HMR (20) while directional nucleation restricts HML silencing (22). In contrast, subtelomeric silencing is rather constrained than restricted. Although numerous factors such as nuclear pore complex components and transcription factors display barrier properties in boundary assays, their physiological role *in vivo* remains to be explored (23). The collective action of chromatin modifying enzymes implements chromatin states that potentially decrease Sir3 affinity for nucleosomes. In addition to the acetylation of H4K16 by the SAS-I complex, acetylation of histone H3 tails by Gcn5 and Elp3, methylation of H3K4 and H3K79, and H4K16ac-dependent incorporation of the H2A.Z histone variant were all proposed to contribute to limit Sir3 spreading at subtelomeres (7). In mutants lacking those enzymes or marks the SIR complex propagates further away from the telomeres (24–27). However the respective contribution of each mechanism and what further limits silencing spreading in those mutants remains unknown.

A key parameter regulating heterochromatin dynamics, function and spatial distribution is the concentration of silencing factors. For instance, increasing Sir3 dosage in budding yeast expands subtelomeric silent domains toward the chromosome core (28) and increases telomere clustering (29,30). However, the dose-dependency of heterochromatin propagation is largely unknown. Intriguingly, Sir3 was recently shown to expend inward chromosomes in G1 arrested cells (31). Here, we discovered that expanded subtelomeric silent domains (ESD) eventually reach a maximum, despite further increases in Sir3 levels. We conducted a meta-analysis and show that ESDs coincide with a pre-existing chromatin landscape as they end at zones of abrupt transition for specific histone marks: increased H3K4me3, H3K36me3 and H3K79me3 and decreased H2AP. Using genetic assays, we identified H3K79me3 as the last bastion of euchromatin protection against the spread of heterochromatin. Finally we show that ESD isolate both structural and functional features providing a new view-point on how to separate subtelomeres from the core chromosome.

## Results

### Saturation of extended silent domains upon SIR3 overexpression

To systemically examine the impact of elevated Sir3, we generated yeast strains stably overexpressing different levels of *SIR3*. We replaced the endogenous *SIR3* promoter with three different promoters, generating strains that produced 1x-Sir3 (WT), 9x-Sir3 (pADH-SIR3), 16x-Sir3 (pTEF-SIR3), and 29x-Sir3 (pGPD-SIR3), as determined by Western blot (FigS1A), and fluorescence quantification of live cells expressing Sir3-GFP (Fig1A, FigS1B). FACS profiles of WT and pGPD-SIR3 strains were largely similar, suggesting that the cell cycle was unaffected by overexpression of Sir3 (SupFigS1D).

**Figure 1:**
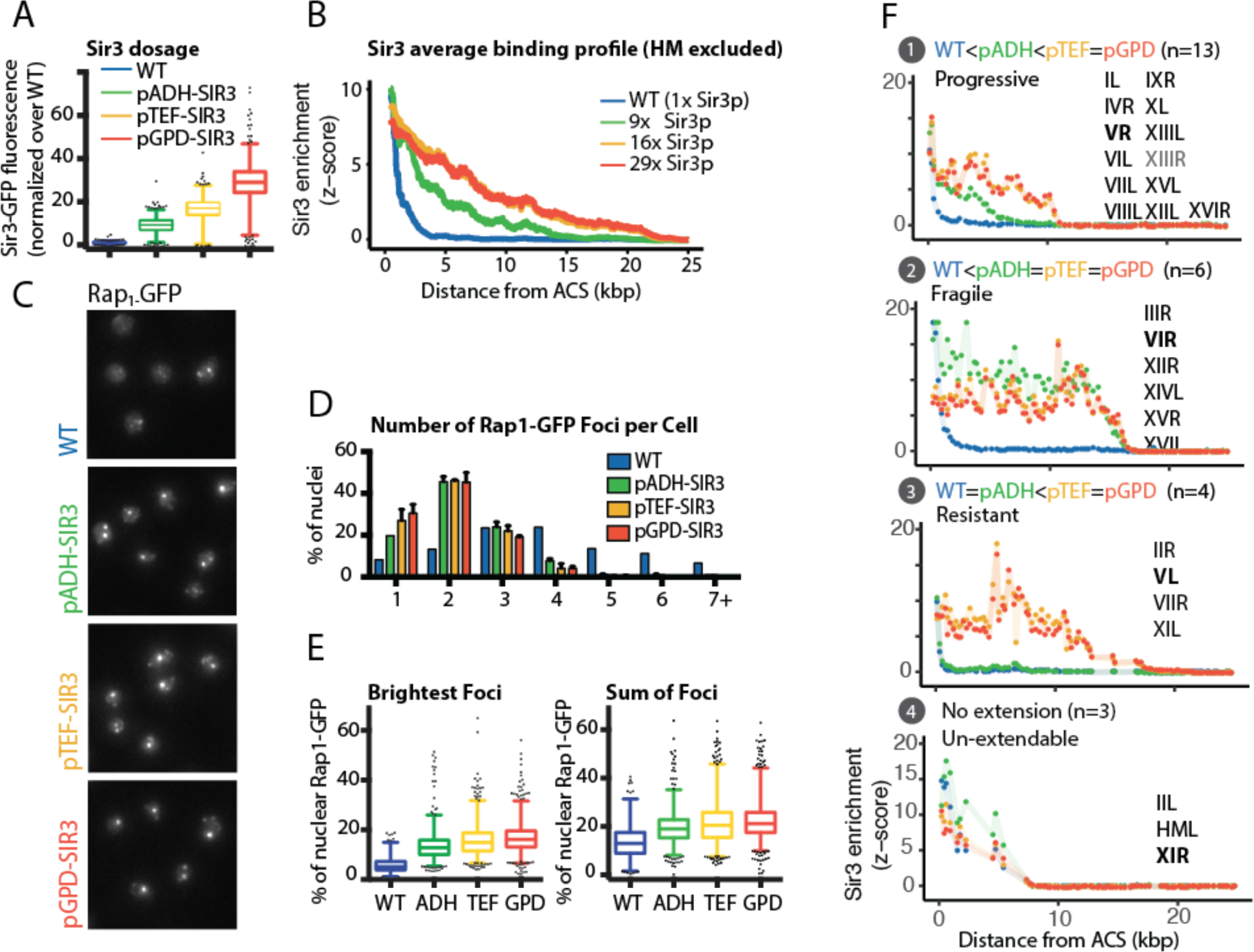
Increasing Sir3 dosage leads to telomere clustering and SIR spreading saturation. (A) Quantification of Sir3 levels by integration of Sir3-GFP signal in strains expressing SIR3-GFP (B) ChIP-chip against Sir3 was carried in strains from A. Moving average of Sir3 binding (block = 1000 bp, window = 10 bp) at telomeres (with the exception of TELIIIL and TELIIIR which contain HM loci) as a function of distance from telomeric X core sequence, the last telomeric element. Enrichment is measured as the standardized IP over Input (See mat meth). (C) Rap1-GFP foci grouping in strain differing for Sir3 levels. Cells were grown in YPD overnight, diluted to OD600nm= 0.2, and imaged at OD600nm= 1. (D) Quantification of Rap1-GFP foci distribution in images from C. (E) Left: Distribution of Rap1-GFP signal attributed to the brightest foci in each nucleus; Right: Distribution of the relative amount of Rap1 measured within foci relative to total nuclear Rap1 signal. (F) Stereotypical examples of Sir3 binding in function of Sir3 dosage, numbers correspond to the subtelomeres constituting each group, in bold is the subtelomere plotted.

To investigate Sir3 spreading, we probed genome-wide Sir3 binding by ChIP-chip. In WT cells, we detected Sir3 binding up to 6-8 kb away from the telomeric repeats, on average, consistent with previous studies (25,32). Elevated levels of Sir3 expanded the distance of Sir3 binding to ~15 kb away from telomeric repeats in 9x-Sir3 cells and up to ~22 kb away in 16x‐ or 29x-Sir3 cells (Fig1B). Sir3 spreading was similar in 16x‐ and 29x-Sir3 cells, but Sir3-GFP nuclear background levels almost doubled in 29x-Sir3 compared to 16x-Sir3 cells (Sup FigS1C), arguing that Sir3 binding to chromatin reached saturation.

In parallel, we monitored telomere foci in function of Sir3 concentration by live microscopy imaging of Rap1-GFP (Fig1C). In wild type cells, telomeres cluster together in 3 to 5 foci located at the nuclear periphery (33). However, upon Sir3 overexpression using the Gal1p promoter, most of the telomeres group together in the center of the nucleus (29). In the range of concentration probed, we observed that telomere clustering increase non-linearly in function of Sir3 levels and saturates at levels between 9-16x Sir3 as we did not detect significant differences between in foci intensity or distribution in cells overexpressing 16x or 29x Sir3p (Holm-Sidak’s multiple comparisons test). Most cells (78%) show at least 3 Rap1-GFP foci in the WT strain while 64-76% of cell show one or two foci in strains overexpressing Sir3 9X or more. Increased foci intensity paralleled the decrease in foci number (Fig1E, Left), consistent with increased telomere grouping in cells overexpressing Sir3. Furthermore, the proportion of nuclear Rap1-GFP within foci increases from 13.6% in WT cells to a maximum of 21.6-22.2 % for Sir3 dosage above 16x (Fig1E, Right). Together, this suggests that not all telomeres are clustered within Rap1-GFP foci in WT cells at a given time, and that elevated Sir3 levels increase the total number of telomeres within clusters, eventually reaching saturation.

Together, these data indicate that increased Sir3 dosage expands Sir3 genome binding and telomere clustering until they reach saturation, between 9-16x Sir3. Some secondary recruitment sites, away from the XCS regions, have been observed previously upon transient Sir3 overexpression(34), creating discontinuous Sir3-bound domains in subtelomeric regions. Interestingly, we observed that the constitutive overexpression of Sir3 submerges those secondary recruitment sites leading to the formation of extended continuous repressive domains.

When looking at individual chromosomes, we observed that Sir3 binding increased at few euchromatic (non subtelomeric) sites upon Sir3 overexpression (FigS1E) but chose not to pursue this further as this was not associated with changes in gene expression in agreement with previous reports (35). Interestingly, individual telomeres showed different behaviours in response to increased Sir3 dosage. We classified telomeres based on their response to Sir3 dosage elevation (Fig1G). “Fragile” subtelomeres (6/26) displayed increased Sir3 spreading and plateaued at 9x Sir3. “Progressive” subtelomeres (13/26) displayed gradually increased Sir3 spreading between 9-16x Sir3 and then plateaued at 16x Sir3. “Resistant” subtelomeres (4/26) displayed increased Sir3 spreading and plateaued at 16x Sir3. Finally, “insensitive” subtelomeres (3/26) did not expand in response to elevated Sir3 levels.

The expanded Sir3 domains showed diverse lengths in all categories, ranging from 7-25kb (HM excluded), independently of chromosomal arm length or middle repeat content (not shown). Thus, extension of Sir3 bound domains exhibits subtelomere specificities but eventually reaches saturation.

### Sir3 spreading extends Silent Domains

Overexpression of Sir3 strongly repressed subtelomeric transcription, as expected. Given that overexpression of the point mutant *sir3-A2Q*, which only binds to telomeric repeats and leads to telomere clustering (Fig2A,B) did not affect global transcription of subtelomeres (FigS2A), repression is attributable to Sir3 binding to chromatin and not clustering of telomeres. However 22 genes showed transcriptional changes common to the over-expression of *SIR3-A2Q* and *SIR3*, *i.e* potentially caused by telomere clustering, including 20 euchromatic genes (FigS2D). Those transcriptional changes could be the consequences of changes in spatial localization or alternatively due to the sequestering of specific factors within the telomere hyper-clusters.

**Figure 2:**
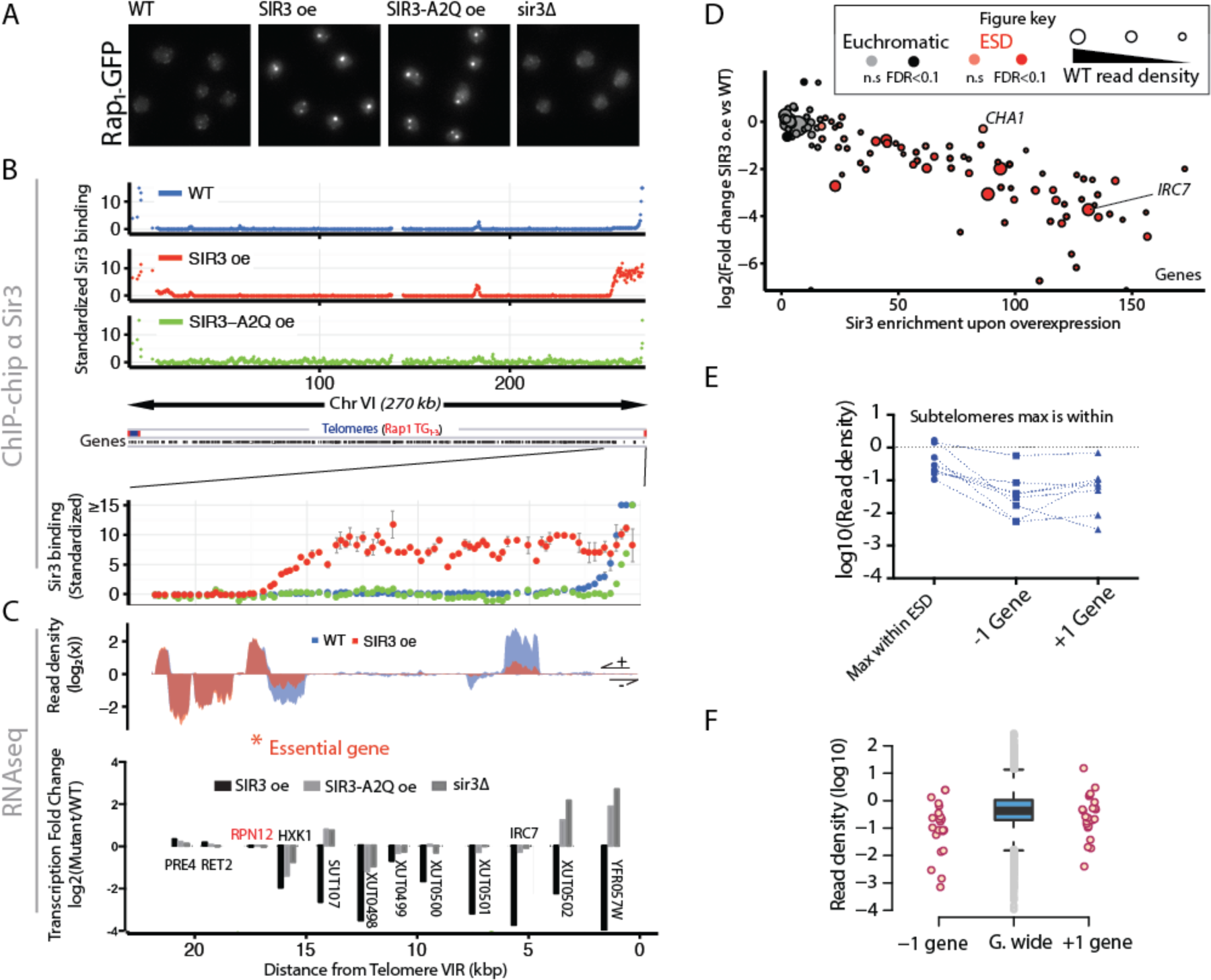
Sir3 extended domains are silenced and restricted to subtelomeres. (A) Representative Rap1-GFP images of exponentially growing strains with different Sir3 amount or expressing the SIR3-A2Q point mutant. (B) Chromosome wide binding of Sir3 in the same strains as in A and blow-up on subtelomere VIR. Enrichment is shown as the standardized IP over Input and scale is thresholded at 15 for visualization purposes. (C) Total RNAseq read density and corresponding transcriptional fold change along subtelomere VIR in indicated exponentially growing (OD~1) strains. (D) Sir3 binding and corresponding transcription changes of subtelomeric genes (Distance from chromosome end <50 kb) upon overexpression of SIR3. Color code indicates if a gene is annotated as within ESD (see math et meth) and shade indicates significance (FDR<0.1) of the detected changes. Read density in WT cells is proportional to the disk area. (E) Exemplification of the 7 subtelomeres at which a gene within E.S.D shows larger transcript amount than the genes located at the end of the domain. (F) Read density of genes located before and after the end of extended silent domains compared to genome wide distribution (central boxplot).

The extension of Sir3-bound domains upon Sir3 overexpression systematically repressed underlying transcripts genome-wide (Fig2C for the Tel6R and 2D genome wide). Repression was largely independent of initial transcript level (Fig2D) and of coding status (e.g. the right subtelomere of chromosome VI, Fig2C, FigS2B). These extended silent domains (ESD) included 100 genes that were not bound by Sir3 in WT cells. The logarithm of transcriptional repression was linearly proportional to the Sir3 binding signal, reflecting the absence of silencing escapers and the correlation between RNA-seq and ChIP-chip (Fig2D). Analysis of reads mapping to multiple loci indicated that entire gene families, characteristic of subtelomeres and Y’ elements, are repressed upon Sir3 overexpression (FigS2C, S2E), suggesting that the subtelomeric regions devoid of chip probes are collectively silenced within ESDs.

Most genes within ESDs are not highly transcribed in WT cells, suggesting that Sir3 spreading might be limited by transcription. However, highly expressed genes like *IRC7* (Fig2C) and *DLD3* (Fig3B) were not excluded from ESDs and were strongly repressed upon *SIR3* overexpression. Both genes belong to the decile of most expressed genes and to the top 20% of most frequently transcribed genes in wild-type cells (36). Similarly, at 7 subtelomeres at least one gene within the ESD had higher read density than the gene adjacent to the ESD (Fig2E). Furthermore gene found immediately before and after the end of ESDs showed similar transcript levels (Fig2F). Therefore, transcriptional activity *per se* appears not sufficient to limit Sir3 spreading when Sir3 is over-abundant.

**Figure 3:**
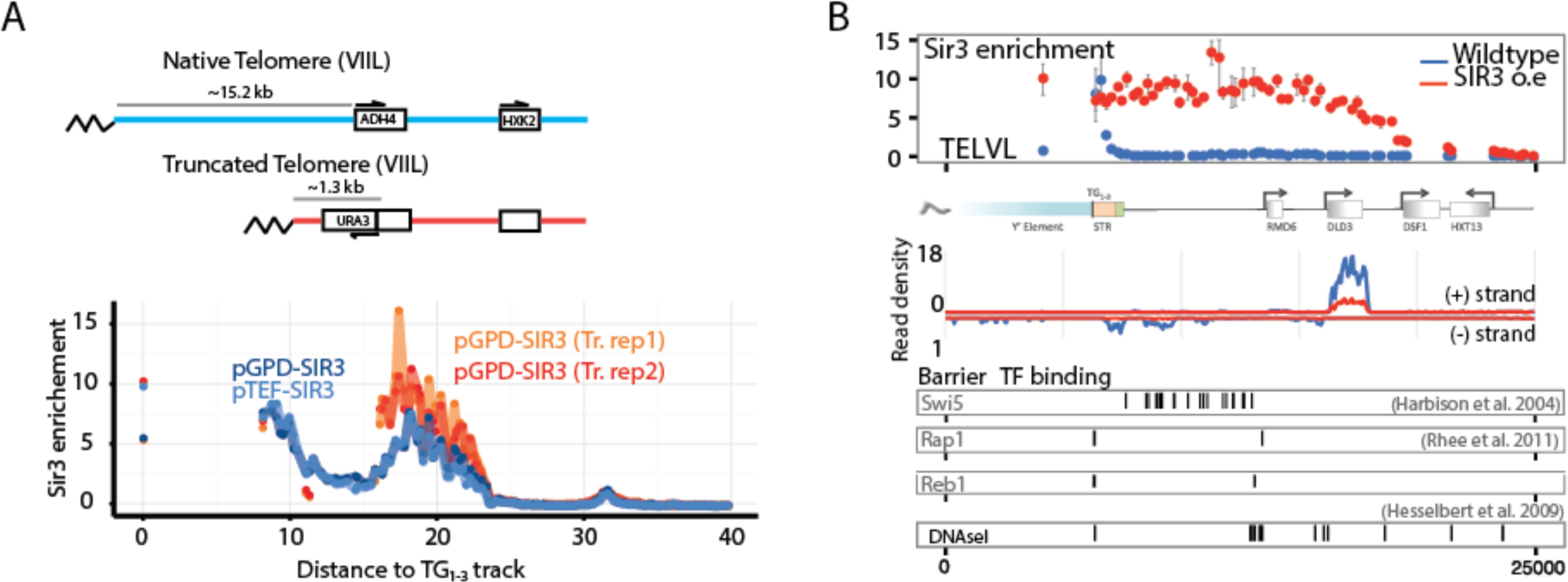
End of extended silent domains is defined locally and independently of transcriptional activity. (A) Sir3 binding at native and truncated TELVIIL, x coordinates correspond the native telomere VIIL (B) Sir3 binding at TELVL in WT and Sir3 overexpressing (pGPD-SIR3) strains. Transcription factor binding and DNAseI hypersensitive sites along TELVL are shown.

The limitation of Sir3 spreading could be the consequence of the counter selection of cells silencing essential genes as ESD did not contain any essential genes and 3 ESD ended right before three essential genes. However, we do not favour this hypothesis for two main reasons. First, we did not detect significant decreases in mRNA levels for these genes upon Sir3 overexpression; second, none of these genes show any haplo-insufficiency phenotype (37) arguing that a weak repression would not be counterselected. We thus conclude that extended Sir3 spreading led to efficient gene silencing of the underlying genes and that gene activity did not account for the end of ESDs.

### ESDs are not limited by distance from the telomere or by previously reported barrier TF elements

To test whether the distance from the telomere limits Sir3 spreading, we compared Sir3 spreading at a WT telomere VIIL versus a 15 kb truncated version (Fig3A). Strikingly, Sir3 binding ended within the *HXK2* promoter in both cases, with a somewhat sharper decline rate in the truncated strains. These data suggest that the Sir3 boundary is either defined relative to the core of the chromosome or is a local feature. Focusing on silent domain ends, we quantified the slope of Sir3 binding at each subtelomere in the strains overexpressing *SIR3*, whenever it was possible (24/32 subtelomeres). We found that the slope at the end of a silent domain did not correlate with the distance from the telomere (*i.e* nucleation point) and found no correlation with the groups defined based on sensitivity to Sir3 dosage (FigS3A). Thus, when Sir3 is in excess, the delineation of the ESD does not depend on the distance from the nucleation site. To investigate whether other DNA sequence-specific barrier elements confine Sir3 ESDs within subtelomeres, we evaluated the available binding data of 10 transcription factors with proposed barrier activity (Adr1, Gcn4, Rgt1, Hsf1, Sfp1, Reb1, Abf1, Leu3, Swi5: (38), Rap1: (39),Tbf1: (40)). At 12 subtelomeres, we identified bound TF sites at genes corresponding to the ESD limit (FigS3C). However, each of these TFs was also bound elsewhere within the ESD (Fig3B) indicating that they are not sufficient to limit the spreading of Sir3. Interestingly, only the three subtelomeres categorized as insensitive to Sir3 levels (group 4), contained known barrier elements flanking Sir3 bound domains: a Leucine tRNA at subtelomere IIL, a previously identified barrier sequence homologous to the left barrier of HML (41) at the subtelomere XIR and the I silencer at subtelomere IIIL (FigS3B).

Thus apart for these three cases, none of the previously identified barrier elements that we could probe was sufficient to block ESD.

### Sir2 does not limit most extended silent domains

We considered that Sir3 spreading might be limited by the capacity of Sir2 within the SIR complex to deacetylate H4K16. We first monitored the genome-wide occupancy of Sir3 in strains overexpressing Sir2. We found that Sir2 overexpression had a limited impact on Sir3 spreading that was weaker than the 9x Sir3 overexpression (FigS4A). Sir3 spreading in cells co-overexpressing Sir2 and Sir3 or Sir3 alone were identical at most subtelomeres (19/26). In the remaining 7 cases, Sir3 spreading increased with co-overexpression of Sir2 (FigS4B), thus slightly extending the average profile of Sir3 binding. It is noteworthy that the further extended silent domains remained devoid of essential and tRNA genes. Thus, Sir2 activity does not generally limit the spread of heterochromatin, even when Sir3 is in excess.

### ESDs encompasses known domains of Sir3 extension

We undertook to compare how Sir3 bound domains extend upon overexpression to known situation of Sir3 binding extension: Sir3 spreading in H3 tail mutants (25) and in cells blocked in G1 by alpha-factor treatment (31).

As shown on figure 4B, ESDs encompass the domains bound by Sir3 in H3 tail mutants or in G1 blocked cells. Interestingly, in the H3 tail mutant, Sir3 bound domains increased only at half subtelomeres but in these cases Sir3 binding profiles were very similar to those observed upon overexpression of Sir3 (Fig4B and FigS4B). In contrast, Sir3 binding in G1 blocked cells appeared to cover domains identical to E.S.D but with a binding signature qualitatively different, as only low magnitude binding signal is observed in the extended Sir3 binding domains.

**Figure 4:**
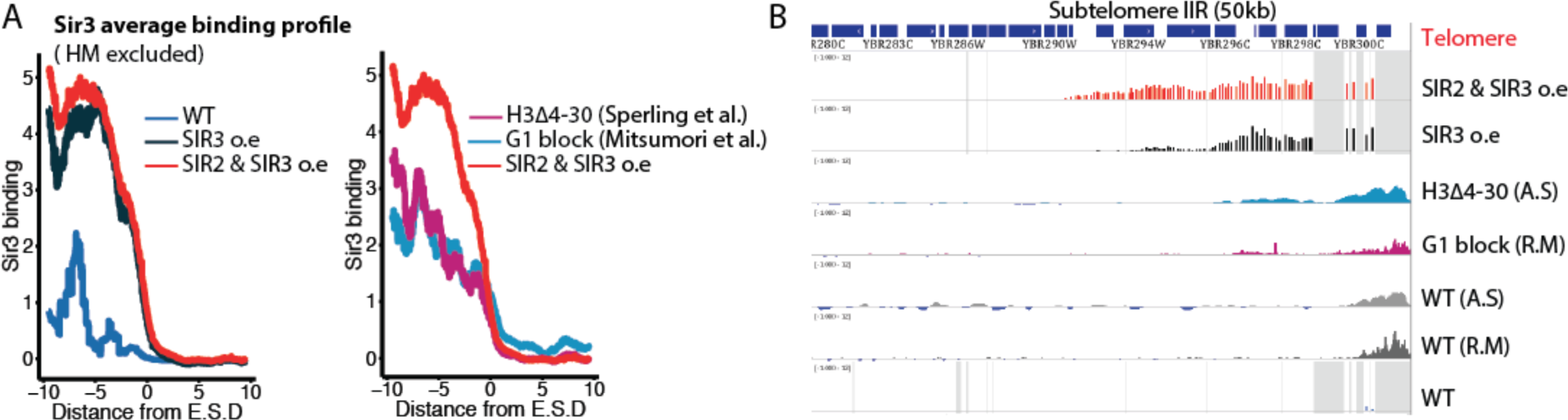
ESDs encompasses known domains of Sir3 extension. (A) Moving average of Sir3 binding (block = 1000 bp, window = 10) at the end of ESD in the indicated conditions or mutants. (B) Genome browser visualization of Sir3 binding at subtelomere IIR, all data are shown as Z-score with a lower bound of −1 and an upper bound of 12.

Together, these data show that Sir3 bound domains in G1 blocked cells or in H3Δ4-30 are contained within E.S.D, although Sir3 is not overexpressed in these conditions. This suggests that the same determinants of Sir3 restriction are at play in all these contexts. Finally, the similarities of Sir3-bound domains in those conditions further suggest that ESDs correspond to chromosomal features that exist independently of Sir3 dosage.

### ESDs coincide with a pre-existing chromatin landscape

To identify potential chromatin determinants of ESDs, we analysed the genome-wide distribution of 26 histone marks or variants (42). We first computed the correlation between Sir3 binding signals and histone modifications at subtelomeres (Fig5A). Consistent with previous results, we uncovered the expected anti-correlation between Sir3 binding and H4K16 acetylation in WT cells. Generally, in wild-type cells Sir3-bound nucleosomes were depleted of most histone marks, with the exception of H4R3 methylation and H2A phosphorylation, which were enriched within silent domains, as reported earlier (43,44). Surprisingly, we observed that Sir3 binding signal is better correlated with several histone marks in all conditions corresponding to extended binding of Sir3 (H3 tail mutants, G1 blocked cells and Sir3 and Sir2/Sir3 overexpression) than in asynchronous wild-type cells. Namely, Sir3 binding signal better correlated with histone H3 methylation and histone H2A phosphorylation (Fig5A, B), the highest correlation values being with Sir3 binding signal in cells co-overexpressing *SIR2* and *SIR3*. Interestingly, while Sir3 binding in G1 blocked cells remains negatively correlated with H4K16 acetylation status, this anti-correlation was much weaker in H3 tail mutants and even lower in strains overexpressing Sir3. This last observation suggests that H4K16 acetylation might limit Sir3 binding in G1 blocked cells but not in H3 tail mutants or upon Sir3 overexpression.

**Figure 5:**
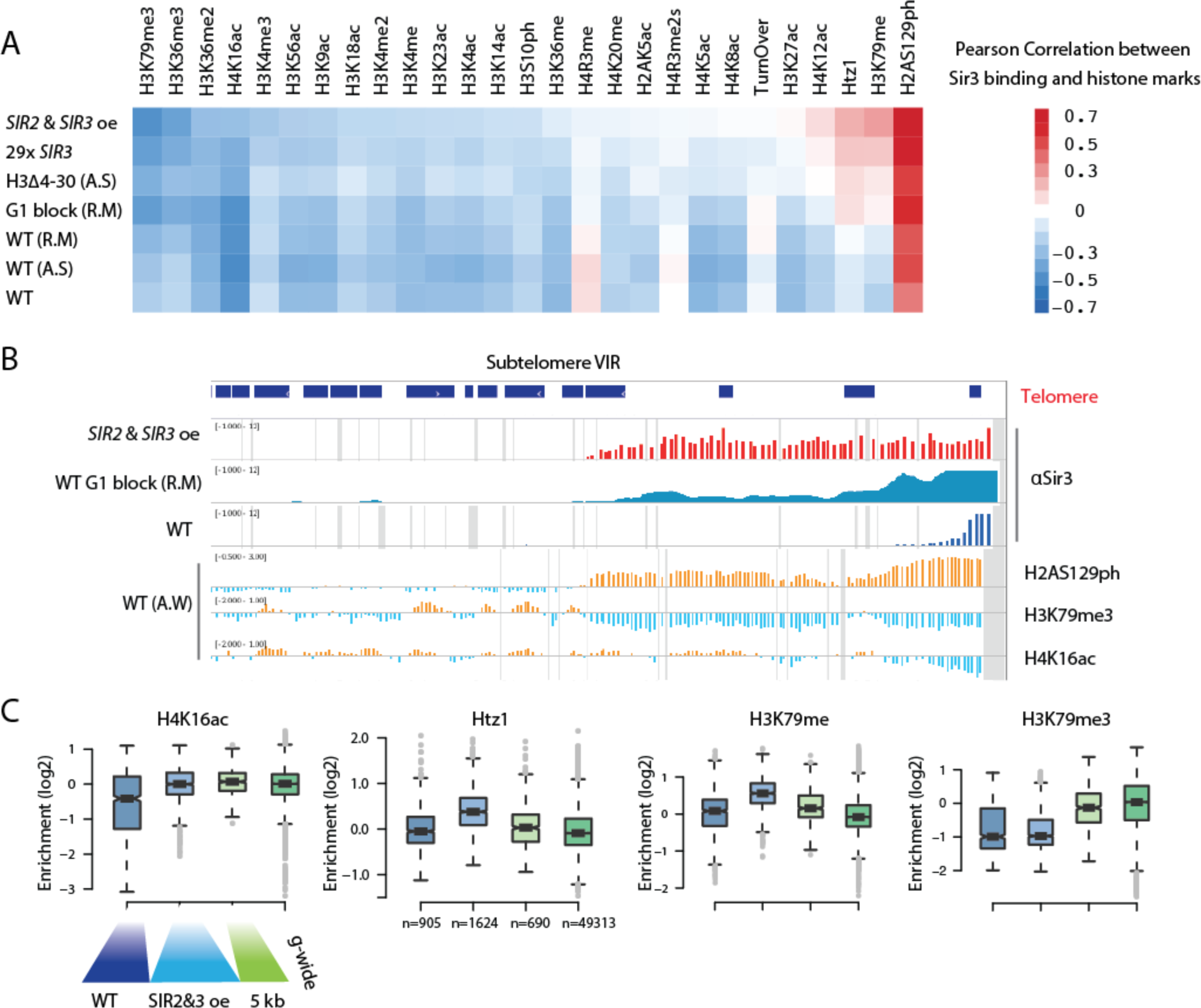
Extension of silent domains predicts major subtelomeric chromatin transitions. (A) Pearson correlation matrix between Sir3 binding and histone marks, 29x Sir3 corresponds to yAT1254 and *SIR2* & *SIR3* oe to yAT1668, external dataset sources are indicated. (B) Genome browser visualization of Sir3 binding in WT, *pGPD-SIR2 pGPD-SIR3* strains, in G1 blocked cells, in H3Δ4-30 mutants and selected histone modification (from Weiner et al.2015) in WT strains at TELVIR. (C) Distribution of selected histone marks relative to H3 (data from Weiner et al. 2015) along wild type silenced domains and within the contiguous subtelomeric domains accessible to Sir3 upon overexpression. As a control, the distribution of those marks within the 5 kb contiguous to the end of extended silent domains as well as the genome wide distribution of those marks is shown.

Surprisingly, H3K79me and the histone variant H2A.Z, previously reported as antagonistic to SIR spreading were depleted from wild type silent domains but enriched within the ESDs and at background levels past ESDs (Fig5AC). This suggests that these chromatin features do not efficiently block Sir3 spreading when Sir3 is over-abundant or in G1 phase.

In contrast, H3K4me3, H3K36me3 and H3K79me3 were depleted until the ESD ends (Fig5C, S5). We reasoned that the longer intergenes present within subtelomeres might bias our analysis, artificially leading to the depletion of marks associated to gene bodies. To control for this potential artifact, we also separated promoter nucleosomes (−3, −2, −1) from gene body nucleosomes and obtained essentially the same results (not-shown).

Thus, subtelomeric chromatin landscape exhibits more similarities with Sir3 binding when it is extended than Sir3 bound region in wild-type asynchronous population.

### H3K79 methylation is essential to protect euchromatin from the spread of silencing

In a complementary approach, we focused on Sir3 binding signal at the ends of ESDs. We classified each subtelomere according to the area under the curve (AUC), corresponding to the logistic-like fit of the Sir3 binding signal in ESDs (see material and methods). At the ten telomeres showing the sharpest change in Sir3 binding at the end of the ESD (highest AUC), several histone marks also displayed sharp changes, particularly H3K79me3 (Fig6A) and H2AS129P (FigS6A). In contrast, the bottom ten subtelomeres that showed gradual changes in Sir3 binding at ESD ends showed rather smooth changes for these marks (Fig 6A and S6A). Thus, ESDs correlate with a pre-existing chromatin landscape defined by specific histone modifications: low levels of H3K79me3 and H3K36me3 and high levels of H2AP.

**Figure 6:**
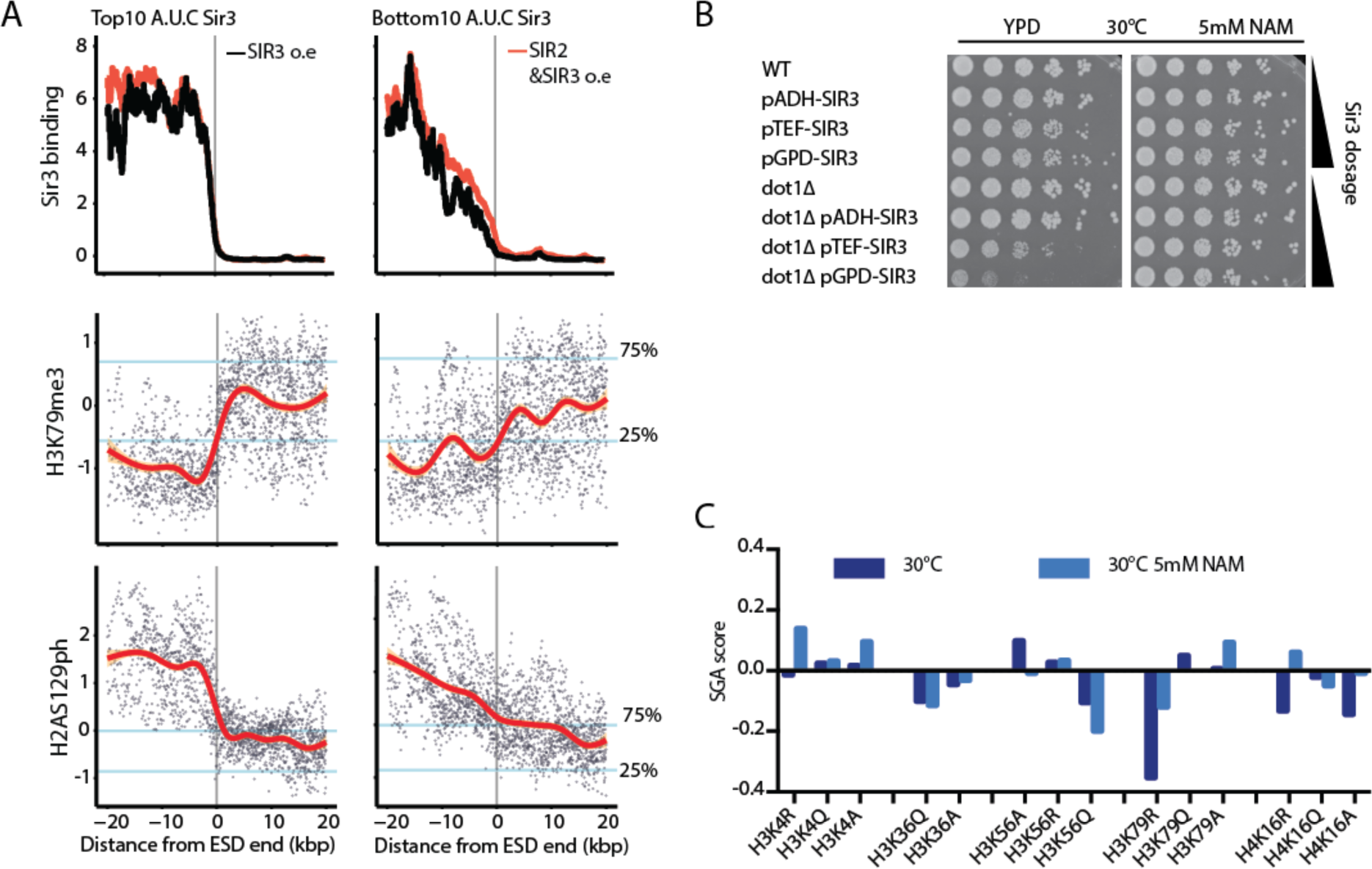
H3K79 methylation is key to sustain viability upon Sir3 overexpression. (A) Moving average of Sir3 binding at telomeres (10kb windows, 10 bp step). The top and bottom 10 telomeres with regards to Sir3 signal in strains overexpressing Sir2 and Sir3 were ploted separately. H3K79me3 were obtained from Weiner et al. 2015. Blue lines indicate genome wide lower and higher quartiles for each mark. Red line corresponds to the local smoothing of histone modification data. (B) Dilution assay to probe viability of *dot1* mutants upon overexpression of Sir3. Cells were constantly grown in presence of 5mM NAM prior to this assay. Cells were grown overnight, and 0.5 O.D of cells were plated in 5x serial dilutions on YPD or YPD 5mM NAM. (C) Growth score of selected histone point mutants on galactose plates (Sir3 inducing conditions) with or without NAM compared to glucose plates (Sir3 dosage is WT).

As H3K79 methylation has been shown to impair Sir3 binding *in vitro* (45,46), this mark represented a good candidate to limit ESDs. To test this hypothesis, we overexpressed Sir3 in this absence of Dot1, the only methyltransferase responsible for H3K79 mono-, di‐ and trimethylation. We found that the *GPD-SIR3 dot1Δ* strains were sick and generated suppressors upon streaking. To avoid artifacts due to these potential escapers, we selected *GPD-SIR3 dot1Δ* clones in the presence of 5 mM nicotinamide (NAM), which efficiently inhibits Sir2 and thus silencing (47). After the selection of positive clones, we assessed the growth of these mutants on medium without NAM, allowing silencing to initiate (48). We found that Dot1 is required for viability when Sir3 is overexpressed >9x (Fig 6B). In contrast, the H3K4 methyltransferase Set1, H3K36 methyltransferase Set2, and histone deacetylase Rpd3 were all dispensable to maintain viability in presence of high Sir3 dosage (FigS6B). Thus, among the chromatin modifications best anti-correlated with limiting ESD, only H3K79 methylation appears essential to restrict the ectopic spread of silencing. Interestingly, the lethality of Sir3 overexpression in *dot1* null was only appreciable at Sir3 amounts above 9x, and was fully rescued by 5 mM NAM treatment (Fig5E). Importantly, a *dot1Δ* strain overexpressing the spreading-defective *Sir3-A2Q* point mutant was viable, further supporting that the requirement for Dot1 is to restrict extended spreading of Sir3 (FigS6B). In addition, co-overexpression of *DOT1* and *SIR3* leads to loss of silencing, showing that H3K79 methylation prevails on Sir3 binding (FigS6).

In a complementary approach we set a genetic screen based on the Synthetic Gene Array to identify H3 and H4 histone residues involved in the limitation of Sir3 spreading. We crossed the collection of histone point mutants(49) with a query strain bearing an additional copy of *SIR3* driven by the inducible GAL promoter. We then monitored colony size in glucose (endogenous Sir3 levels) versus galactose medium (high Sir3 levels) in presence or absence of NAM and listed mutants that had defects above our selected threshold. Interestingly, H3K79R was the sole histone point mutant having growth defects that could be rescued by 5 mM NAM treatment (Fig6C, S6E) indicating that this residue plays a key role in limiting Sir3 spreading. H4K16R mutant had non-significant growth defect (Fig6C, S6E), consistent with a rather subtle influence of H4K16 on the maximal extent of silencing upon *SIR3* overexpression. Thus our data strongly suggest that H3K79me3 actively restricts the ectopic spreading of silent chromatin beyond subtelomeric regions thus protecting euchromatin.

### ESDs coincide with discrete domains that segregate subtelomeric features

We have identified discrete subtelomeric domains corresponding to the maximal extension of Sir3 bound domains. We next sought to identify regulators of genes found in these domains by screening a compendium of over 700 mutants (50). We classified subtelomeric genes into four different groups (1) genes or pseudo-genes associated with middle repeat elements (telomeric), (2) genes bound by Sir3 in WT cells, (3) genes bound by Sir3 only upon Sir3 overexpression and (4) genes bound Sir3 only upon co-overexpression of Sir2/Sir3. Groups 3 and 4 correspond to ESDs. We also considered the group of genes located within 10kb from the end of ESDs and located between 10-20kb from ESD ends.

For each mutant, we tested if the proportion of differentially expressed genes (|log2(FC)|>2) within a subtelomeric domain was higher than expected by chance, considering the effect of the mutation on the genome. We identified genes whose mutation affects specific subtelomeric subdomains (Fig7A). As expected, deletion of any of the SIR had localized effects within the ‘telomeric’ and ‘WT Sir3 bound’ domains.

**Figure 7:**
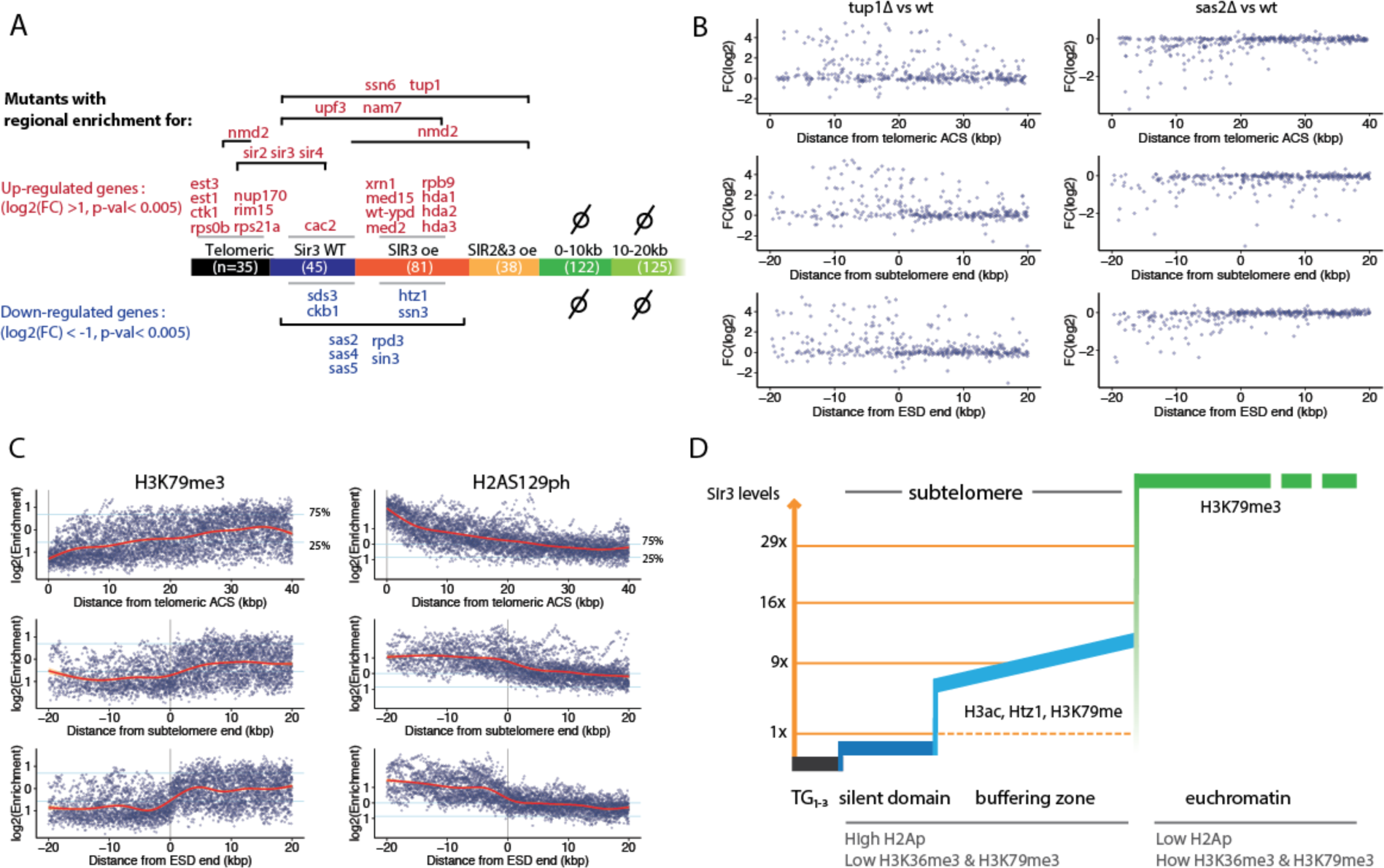
Extension of silent domains reveals new aspects of subtelomeric structuration. (A) Localized effects of mutations affecting subtelomeric transcription. The different subtelomeric subdomains are defined according to Sir3 binding. The number of genes within each domain is into brackets. Mutant names are positioned according to the domain(s) within which the proportion of genes up or down-regulated (log(FC)>1 or < −1) is significantly elevated (Hyper-geometric law, with Bonferonni correction n=703). (B) Transcription changes of subtelomeric genes in *tup1* and sas2 mutants (data obtained from Kemmeren et al. 2014) relative to different subtelomeric viewpoints. Coordinates of subtelomere ends were obtained from Yue et al. 2016. (C) Distribution of H3K79me3 and H2AS129ph (obtained from Weiner et al. 2015) relative to different subtelomeric viewpoints. Blue lines indicate genome wide lower and higher quartiles for each mark. Red line corresponds to the local smoothing of histone modification data.(D) Model depicting how extending silent domains enables to uncover consistent subtelomeric domains delimited by chromatin mark transitions.

Our analysis retrieved mutants previously known to affect subtelomeric transcription. Telomerase components and the nucleoporin *NUP170* (51) up regulates specifically the most telomeric group of genes in our analysis, while the mediator complex tails proteins Med2, Med15 and Gal11 (52,53), the Hda1/2/3 complex, and the general repressors Tup1/Cyc8 specifically affect genes located within ESD (Fig7A). Importantly, the localized enrichment for downregulated genes in *rpd3* or *sas2/4/5* mutants did not extend beyond the ESDs. This enrichment for downregulated genes is likely the consequence of increased spreading of the SIRs in those mutants. Therefore, SIR dependent silencing in those mutants seems not to extend beyond ESD, further reinforcing the notion that ESD represent the maximal extend of SIR dependent silencing. Significantly, this analysis reveals that all the mutants specifically affecting subtelomeric transcription impact domains that are contained within ESD. This suggests that ESD captures well those properties of subtelomeres.

Lastly we wished to probe how the ESD point of view segregates subtelomeric properties as compared to other definitions of subtelomeres: distance from telomeres, from the first essential gene or based on synteny conservation across evolution (54). ESD often overlapped with the core chromosome as defined by synteny (FigS7A). Interestingly, ESD better segregates transcriptional changes associated to *TUP1* or *SAS2* deletion (Fig7B, S7B) or histones marks such as H3K79me3 or H2AS129ph (Fig7C) than the other definitions of subtelomere do. In the case of transcriptional changes associated to tup1 deletion, the sharp p-value transition of proportionally enriched domains coincides with ESD ends (FigS7B). Thus probing the maximal extent of silencing domains reveals discrete subtelomeric domains delimited by histone mark transition zones and provides a new definition of subtelomeres (Fig7D).

## Discussion

The Sir complex has been a model for chromatin complex propagation and gene silencing for decades. Pioneer studies demonstrated that increasing the dose of Sir3 extends silenced domains at subtelomeres (55,56), a property common to several heterochromatin complexes. However there has been controversy on the generality of this finding at natural telomeres (56), and the details of this process remain unclear. Here, we systematically studied the impact of increasing Sir2 and Sir3 dosage on the propagation of the SIR complex and on genome-wide transcription.

Gradual overexpression of Sir3 revealed that the concomitant spreading of Sir3 over subtelomeres reaches saturation at Sir3 levels between 9 and 16x. Unexpectedly, the responses to increase in Sir3 levels were not continuous at all subtelomeres, and the concentration of Sir3 required for maximal spreading varied. The extent of spreading varied up to 30 kb in-between subtelomeres, independently of middle repeat elements or chromosomal arm length. A few telomeres are constrained by punctual elements and insensitive to elevated Sir3. Current models to explain the restriction of heterochromatin spreading include either negotiable or fixed borders (57). Only fixed borders are expected to be independent of silencing factor concentration. Thus, our results suggest that the saturation of silent domain expansion likely correspond to the reaching of fixed borders along most subtelomeres.

Sir3 spreading, and in turn silencing, were largely independent of the underlying gene, sequence or subtelomere under consideration. This suggests that spreading efficiency is mostly dictated by the ability of Sir3 to associate with chromatin. Accordingly, the domains covered by Sir3 upon overexpression shared similar chromatin marks, suggesting that the chromatin landscape dictates the extent of Sir3 spreading.

Our data indicate that the methylation of H3K79 by Dot1 is required for viability when Sir3 is overexpressed. Our work reveals the maximal subtelomeric domains accessible to Sir complex spreading in viable cells, and uncovers previously unknown, discrete subtelomeric domains.

### Different categories of Sir chromatin antagonism

The extent of spreading of the Sir complex at subtelomeres depends on nucleation element strength, chromatin modifying enzymes and Sir concentration (7,11). Ultimately, those parameters influence the affinity of Sir3 for chromatin. Many studies have investigated the consequences of modulating chromatin-modifying enzymes, however we chose to evaluate Sir3 concentration. Upon overexpression, Sir3 binding extends within regions that contain or are enriched for chromatin marks previously reported as antagonistic to its spreading, such as histone variant H2A.Z (27,58,59) and monomethylated H3K79 (45). Although our data suggests the existence of fixed borders, our search for punctual border elements retrieved convincing candidates only at the three subtelomeres for which the extension was already limited in wild-type cells. Furthermore, native binding sites for transcription factors that block silencing when tethered to chromatin (23) were not efficient barriers to Sir3 spreading. Consequently, our work indicates that histone tail acetylation, H2A.Z, and specific transcription factors likely buffer the spread of the SIR rather than block it. The finding that Sir3 binding in G1 blocked cells is better correlated with chromatin landscape than Sir3 binding in asynchronously growing cells suggests that Sir3 might be actively contributing to subtelomeric chromatin landscape at loci previously thought to be unbound by Sir3 when probed on mixed populations of cycling cells.

### End of extended silent domains: the specific role of Dot1

We observed that the end of extended silent domains coincide with a major histone mark transition zone, characterized by the enrichment of H3K4me3, H3K36me3 and H3K79me3. Deletion of *SET1* or *SET2,* the genes encoding the enzymes responsible for the H3K4me3 and H3K36me3, respectively, had no impact on cell growth upon Sir3 overexpression. However deletion of *DOT1*, which encodes the only H3K79 methyltransferase in budding yeast, was lethal in this condition. Furthermore, H3K79R was the sole histone point mutant showing growth defects upon overexpression that could be rescued by inhibiting Sir2 in our genetic screen.

Dot1 is a conserved enzyme, responsible for mono-, di‐ and trimethylation of H3K79 (60). Crystal structure data indicate that K79 methylation would block hydrogen bond formation between K79 and the BAH of Sir3, thereby decreasing Sir3 affinity to nucleosomes (61,62). *In vitro,* all three H3K79 methyl marks abolish the binding of Sir3 to H3 peptides (45,46) and reduced Sir3 affinity for reconstituted nucleosomes (62–64). In vivo, Sir3 was found associated with H3K79 mono and di methylation at active subtelomeric genes (62). However, other studies showed that Sir3 spreading and silencing establishment accelerates in absence of Dot1 indicating that H3K79 methylation prevents Sir3 binding (48,65).

Here we show that upon overexpression, Sir3 spreads over domains enriched for H3K79me, implying that this mark does not block Sir3 spreading *in vivo*, in agreement with previous reports (62). H3K79me3 is mutually exclusive with H3K79me1 (66), so we hypothesized that loss of H3K79me3 is responsible for the deadly spreading of elevated Sir3 in the absence of Dot1. Accordingly *bre1* mutants - that lacks H3K79me3 and H3K4me3 methylation and has increased H3K79me1 (67) - behave similarly in our systematic screen as *dot1*, albeit with a weaker phenotype. Both mutants were synthetic sick with *SIR3* overexpression, a phenotype that was rescued by inhibiting Sir2 (not shown). Our data thus suggests that H3K79me3 observed at the boundary of extended silent domains block the spreading of Sir3, and thus protects euchromatin from heterochromatin.

In agreement with this, co-overexpression of *DOT1* and *SIR3* led to loss of silencing. While we could not differentiate between ESD of WT strains overexpressing 16x and 29x Sir3, these strains exhibited clear differences in the absence of Dot1. This suggests that the ESD saturation with increasing Sir3 depends on Dot1 activity.

We propose that H3K79me3 actively restricts silencing to subtelomeric region thus protecting euchromatin. As the occupancy of this mark is independent of transcription rate (66), this offers the attracting possibility of preventing heterochromatin spreading independently of transcription.

### Subtelomeric specificities

In most organisms, specific features of chromosome ends extend beyond telomeres, within domains generally referred to as subtelomeres (4). In budding yeast, several viewpoints enable the identification of diverse subtelomeric features. A recent study comparing the conservation of synteny among closely related yeast species enabled a precise definition of budding yeast subtelomeres showing a faster evolution than the core genome due to higher CNV accumulation, rampant ectopic reshuffling and rapid functional divergence (54).

The chromatin landscape also exhibits specific features within domains located proximal to chromosome ends (68–70). The first is undoubtedly the presence of heterochromatin, which has a unique signature in terms of histone marks. However specific properties associated with chromosome ends often extend beyond heterochromatic domains (68,69). At most *S.cerevisae* subtelomeres, Hda1-affected subtelomeric (HAST) domains (70) and Htz1 activated (HZAD) domains (58) lie contiguous to SIR silenced chromatin. In addition, phosphorylation of H2AS129 and monomethylation of H3K79 also extend further away than SIR silenced domains. Interestingly, ESDs reveal subtelomeric domains that possess a consistent chromatin signature. Namely these domains are enriched for H2AP, Htz1 and depleted of methylated histone H3 and show sharp transition at the end of ESDs. Consistently, Htz1 sensitive genes are enriched in these domains. Furthermore, considering the end of ESDs as a boundary between subtelomeres and the core genome segregates genes sensitive to the depletion of chromatin modifiers such as hda1, Tup1/Ssn6 or sas2 better than other definitions of subtelomeres do (FigS7). Thus ESDs coincide with discrete subtelomeric domains isolating structural and functional features.

ESDs coincide with the evolutionary definition of subtelomere based on synteny at 7 out of the 26 subtelomeres that we could analyse (FigS7A) and the syntenic chromosome core is accessible to Sir3 spreading at most telomeres up to 27 kb. We and others recently showed that chromatin states impact on efficiency and outcome of both homologous recombination and nucleotide excision repair (71,72). This raises the question of whether the specific chromatin state associated with subtelomeric domains that we uncovered in this study, contributes to the particular evolution of those regions.

### Contribution of telomere proximity to subtelomeric properties

A central question of the biology of subtelomeres is to what extent the properties of subtelomeres are due to their proximity to telomeres or mere consequence of their gene content. Several studies demonstrated that the SIR complex contributes to the localization of enzymes to subtelomeres. For example, subtelomeric localization of the Okazaki fragment processing protein Dna2 is severely reduced in sir mutants (73). In addition, the kinase Tel1 responsible for H2A phosphorylation in subtelomeric regions is present at telomeres but H2AP levels depend mainly on the integrity of the SIR complex (74). Interestingly Sir3 stabilizes this mark even at regions where Sir3 is not detectable by ChIP in wild-type cells, suggesting that either Sir3 acts remotely, or binds these regions at least transiently in wild-type. Intriguingly, regions enriched for H2AP coincide with ESD, leading to the hypothesis that overexpressing Sir3 stabilizes these transient interactions. Accordingly, profiling of Sir3 binding in G1 arrested cells showed low levels of Sir3 binding within ESDs (31). Thus Sir3 might contribute to shape the chromatin landscape in subtelomeric regions.

## Conclusion

By taking the opposite approach to depletion studies, our work describes the dose dependency of budding yeast heterochromatin. In the presence of a large excess of silencing factors, ectopic nucleation of heterochromatin remains limited and does not impact euchromatic transcription. In contrast, we observed the extension of subtelomeric silent domains and characterized their maximal extension along with the antagonistic factors that have been overcome, such as H2Az or H3K79me. By scanning chromatin properties associated with Sir3 maximal binding, we uncovered major subtelomeric histone mark transition zones that functionally protect euchromatin from the spread of silencing. The long-term contribution of heterochromatin to the peculiar properties of subtelomeres will require further study.

## Material and methods

Media and Growth conditions: Yeast cells were grown on YP with 2% glucose, raffinose or galactose. Unless notified, all the strains used in this study were grown at 30 °C with shaking at 250rpm.

### Yeast transformation protocol

Cells were seeded on liquid medium and grown to 0,8<OD_600_<1,2. 3 ODs (~3x10_7_ yeast cells) of cells were taken and washed with 1X TEL (10mM EDTA pH 8, 100mM Tris pH8, 1M Lithium Acetate), then 3μl of SSDNA (Sigma ref: D9156-5ML), DNA template (0,5μl if plasmid DNA, 5μl of digested plasmid or PCR product), 300μl of 1X TEL and 45% PEG-4000 solution were added. The mix was put 30 min at 30 °C and heat shocked at 42°c for 15 minutes. Lastly, cells were plated on appropriate selective medium.

### Rap1 foci analysis

The image analysis is performed with a slightly modified version of the dedicated tool from (75). These modifications regard the quantification of foci and aim at providing a more accurate estimation of the quantity of fluorescence held inside each focus. The gaussian fitting approach has been replaced by a template matching framework with a bank of 100 symmetric 2D gaussian kernels with standard deviations ranging from 0.5 to 7 pixels. The position of each template is determined as the maximum of normalized cross correlation whereas the most suitable template for a single focus is selected by minimizing the sum of square differences between the gaussian template and the data within a circular mask of radius twice the standard deviation. The foci are then defined as spherical objects with radii of two times the standard deviations of the matched templates. All foci that could not be fitted were considered as a cube of dimension 5*5*5. Variation of the box size did not affect overall results. The foci intensity can thus be measured as the sum of the fluorescence signal inside its sphere. Furthermore, the proportion of intensity from a nucleus held inside each of its foci is also computed.

### Sir3-GFP quantification

Quantification of Sir3-GFP signal was carried using microscopy. Briefly cells were segmented on the basis of trans signal using a modified version of CellX, and the intensity of 30 Z-stacks deconvolved imaged was summed. Deconvolution was carried using Metamorph. For each cell the intensity/pixel was measured and normalized by the WT average.

### Western blots

Protein extracts were prepared from 2 ODs of exponentially growing cultures (O.D ~1) using the post-alkaline extraction method (Kushnirov et al. 2000). For immunoblotting, we used custom-made polyclonal antibodies against Sir3 [1:10000].

### FACS

Cell cycle profiles were obtained on a Accury FACS machine using CYTOX as DNA staining agent and analyzed using FlowJoX.

### SGA screen

Query strain was obtained by transforming strain yAT-1949 with pGAL-SIR3-HPH, integrated within *TRP1*. The query strain was crossed with the collection of histone point mutants (Dai et al. 2012) following the selection steps described in (Tong et al, 2001), with selection media adapted to respective genotypes. Each cross was done in quadruplicate on 1536-format plates. Once double mutants were acquired, they were transferred to one of the following medium: double mutant selection medium (glucose), double mutant selection medium (glucose) + 5 mM NAM, double mutant selection medium (galactose) or double mutant selection medium (galactose) + 5 mM NAM. All strains were grown at 30°C and imaged after 2 days. Image analysis and scoring were done with SGAtools (Wagih et al. 2013), where mutants growing on glucose media served as controls. Only significant changes were considered (p-val< 0.001) and a last significance threshold was chosen to only keep mutants which score’s absolute value was >0.2.

### Dilution Assays

Cells were grown overnight in YPD 5mM NAM before dilution. 5X serial dilutions are shown. Plates were grown for 2-3 days at 30°C unless indicated otherwise.

### Pellet preparation for ChIP

A total of 20 O.D equivalent of exponentially growing cells were fixed in 20 mL with 0.9 % formaldehyde for 15 min at 30°C, quenched with 0.125 M glycine and washed twice in cold TBS 1x pH 7.6. Pellets were suspended in 1mL TBS 1X, centrifuged and frozen in liquid nitrogen for -80°C storage.

### Chromatin immunoprecipitation

All following steps were done at 4°C unless indicated. Pellets were re-suspended in 500 μL of lysis buffer (0.01% SDS, 1.1% TritonX-100, 1.2 mM EDTA pH 8, 16.7 mM Tris pH8, 167 mM NaCl, 0.5 % BSA, 0.02 g.L^−1^ tRNA and 2.5 μL of protease inhibitor from SIGMA P1860) and mechanically lysed by three cycles of 30s, intensity 6ms-1 with 500 μm zirconium/silica beads (Biospec Products) using a Fastprep instrument (MP Biomedicals). Each bead beating cycle was followed by 5 min incubation on ice. The chromatin was fragmented to a mean size of 500 bp by sonication in the Bioruptor XL (Diagenode) for 14 min at high power with 30 s on / 30 s off and centrifuged 5 min at 13 000 rpm. 10 μL were kept to be used as Input DNA. Cleared lysate was incubated overnight with 1 μL of polyclonal antibody anti-Sir3. 50 μL of magnetic beads protein A (NEB) were added to the mixture and incubated for 4h at 4°C. Magnetic beads were washed sequentially with lysis buffer, twice with RIPA buffer (0.1% SDS, 10mM Tris pH7.6, 1mM EDTA pH8, 0,1% sodium deoxycholate and 1% TritonX-100), twice with RIPA buffer supplemented with 300 mM NaCl, twice in LiCl buffer (250 mM LiCl, 0.5% NP40, 0.5 % sodium deoxycholate), with TE 0.2% TritonX-100 and with TE. Input were diluted 10x with elution buffer (50mM Tris, 10mM EDTA pH8, 1%SDS) and beads were re-suspended in 100 μL elution buffer. A reversal cross-linking was performed by heating samples overnight at 65°C. Proteins were digested with proteinase K in presence of glycogen and the remaining DNA was purified on QIAquick PCR purification columns. Finally, samples were treated with RNAse A 30 min at 37°C.

### ChIP-chip preparation and hybridation

Samples used for ChIP-chip have all been analysed by qPCR prior to microarray hybridization. For microarray hybridization 4/5 of the immunoprecipitated DNA and of the DNA from the input were ethanol precipitated and re-suspended in 10μL of water (Gibco). Purified material was amplified, incorporating amino-allyl-dUTP using as described in (75). The size of amplified fragments (~500 bp) was assessed by gel electrophoresis. For each sample 1.5 μg of amplified DNA was coupled either with Cy5 (immunoprecipitated sample) or Cy3 (input sample) and hybridized on 44k yeast whole genome tiling array (Agilent) as described in (75).

### Microarray data acquisition, analysis and visualization

Microarray was imaged using a Agilent DNA microarray scanner and quantified using GenePix Pro6.1 as described in (75).

### Genome wide data analysis

All dataset were lifted over to Saccer3 when required. Histone marks data were obtained from (42). Sir3 binding in H3 tail mutants from (25), nucleosome turnover from (76). Transcriptome data were downloaded from the website supporting the publication (50). Subtelomere definition was obtained from (54). Z-scores were computed using the R scale function.

Downsampling of (Sperling et al. 2009) and (Mitsumori et al. 2016) data to the 44k microarray probes for Fig4B and 5A was done using R, visual inspection of the data confirmed that downsampling was carried without errors. Average telomeric profiles were done by computing the mean of the signal over 10 kb windows separated by 10 bp. The limits of Extended silent domains were computed as the first probes possessing 5 neighboring probes that have Z-score inferior to 1, starting from the telomere. Fitting of the data was done using Matlab fitting toolbox using Bisquare robustess option. The function used is f(x)=K/(1+exp(−r*(−x+t0)))+1, with the following fitting parameters for K,r, and t0: lower bounds: [10 0.0001 1000], Starting point: [10 0.0001 1000], upper bounds: [200 0.01 40000]. Area under the curve was exactly computed on the fitted signal of Sir3 binding in strains overexpressing *SIR2* and *SIR3*, 10kb before the end of silent domains and 5 kb after.

Mutants showing localized effects were identified with using the hypergeometric distribution, function phyper with Bonferroni correction for multiple testing (n=703).

### RNAseq

Total RNA from a 25mL culture of exponentially growing yeasts were extracted using phenol-chloroform. Banks were constructed using the kit SOLiD Total RNA-Seq, with minor modifications: RNA are Zinc fragmented and fragments with size ranging form 100 to 200 nt selected by gel purification. After reverse transcription only fragment of size > 150nt are kept. Paired end (50 + 35) sequencing was done by the Institut Curie plateform. Differential expression was called using EdgeR, with a false discovery rate inferior to 0.1.

## Acknowledgments

We thank T. Rio Frio, S. Baulande and P. Legoix-Né (NGS platform, Institut Curie), A. Lermine (bioinformatics platform, Institut Curie), P.Lebaccon (PICT@Pasteur microscopy platform) for support. We also thank Camille Gautier for help with multiple mapping reads analysis and Marta Kwapisz for RNAseq library preparation. We thank Valérie Borde for help with microarray experiments, David Sitbon, Valérie Borde and Arnaud De Muyt for fruitful discussions, David Sitbon and Ann Ehrenhofer-Murray for critical reading of the manuscript. This work has benefited from the facilities and expertise of the NGS platform of Institut Curie (supported by the Agence Nationale de la Recherche [ANR-10-EQPX-03, ANR10-INBS-09-08] and the Canceropôle Ile-de-France), and of the high-throughput sequencing platform of IMAGIF (<u>http://www.i2bc.paris-saclay.fr</u>). A. Morillon’s lab is supported by the Agence Nationale de la Recherche (DNAlife) and the European Research Council (EpincRNA starting grant, DARK consolidator grant). This work was supported by funding from the Labex DEEP (ANR-11-LABX-0044_DEEP and ANR-10-IDEX-0001-02 PSL) and from the ANR DNA-Life (ANR-15-CE12-0007). AH was supported by fellowships from the ENS PhD program and the FRM (DEP20131128535).

## SupplementaryFigures

**Supplementary Figure 1:**
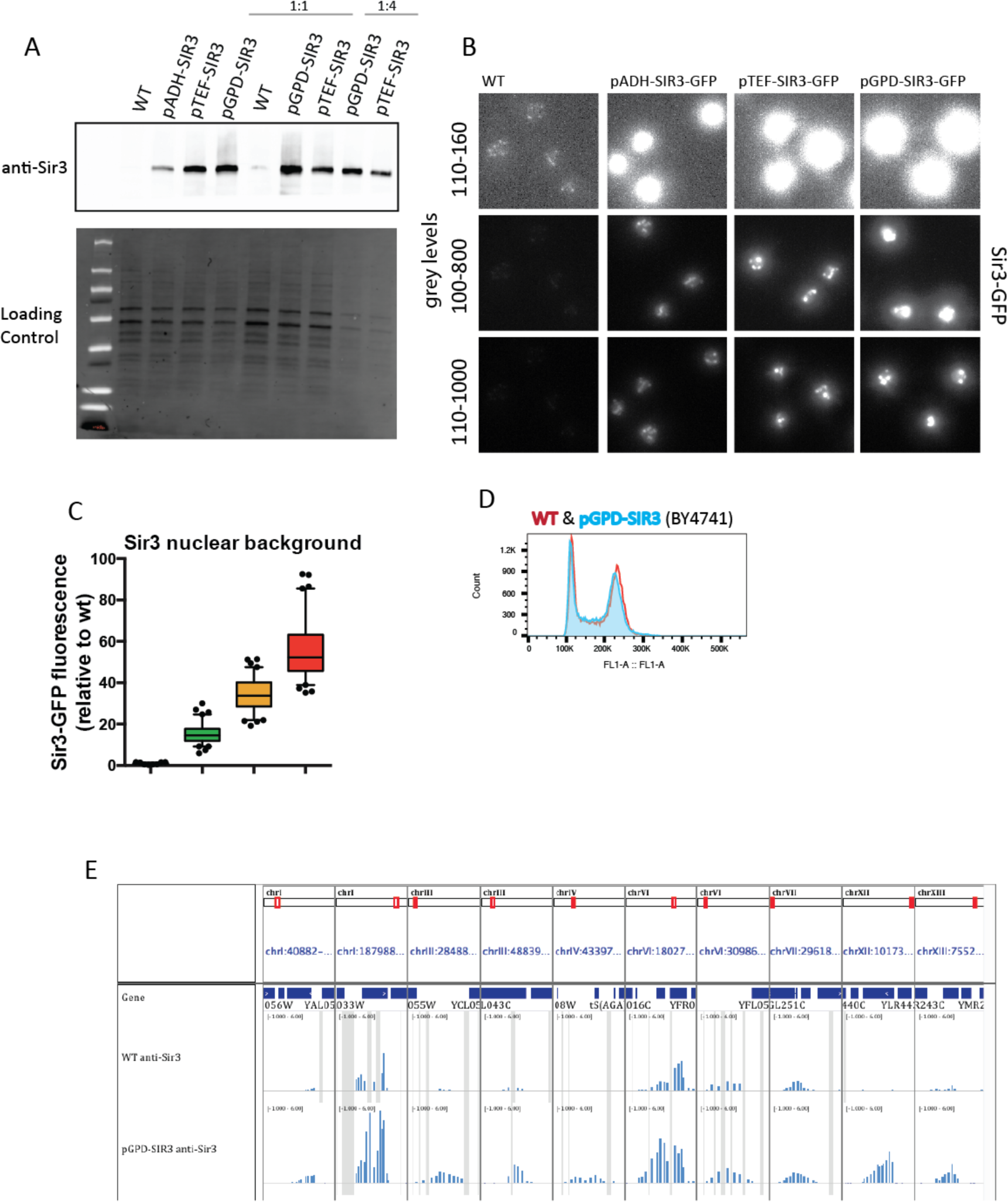
(A) Western Blot anti-Sir3 in the strains used in Figure 1 for ChIP-chip. (B) Representative examples of Sir3 fluorescence in strains overexpressing Sir3-GFP. (C) Quantification of Sir3-GFP nuclear background. (D) FACS profile of exponentially growing WT and *pGPD-SIR3* strains. (E) Representative images of loci bound by Sir3 within euchromatin.

**Supplementary Figure 2:**
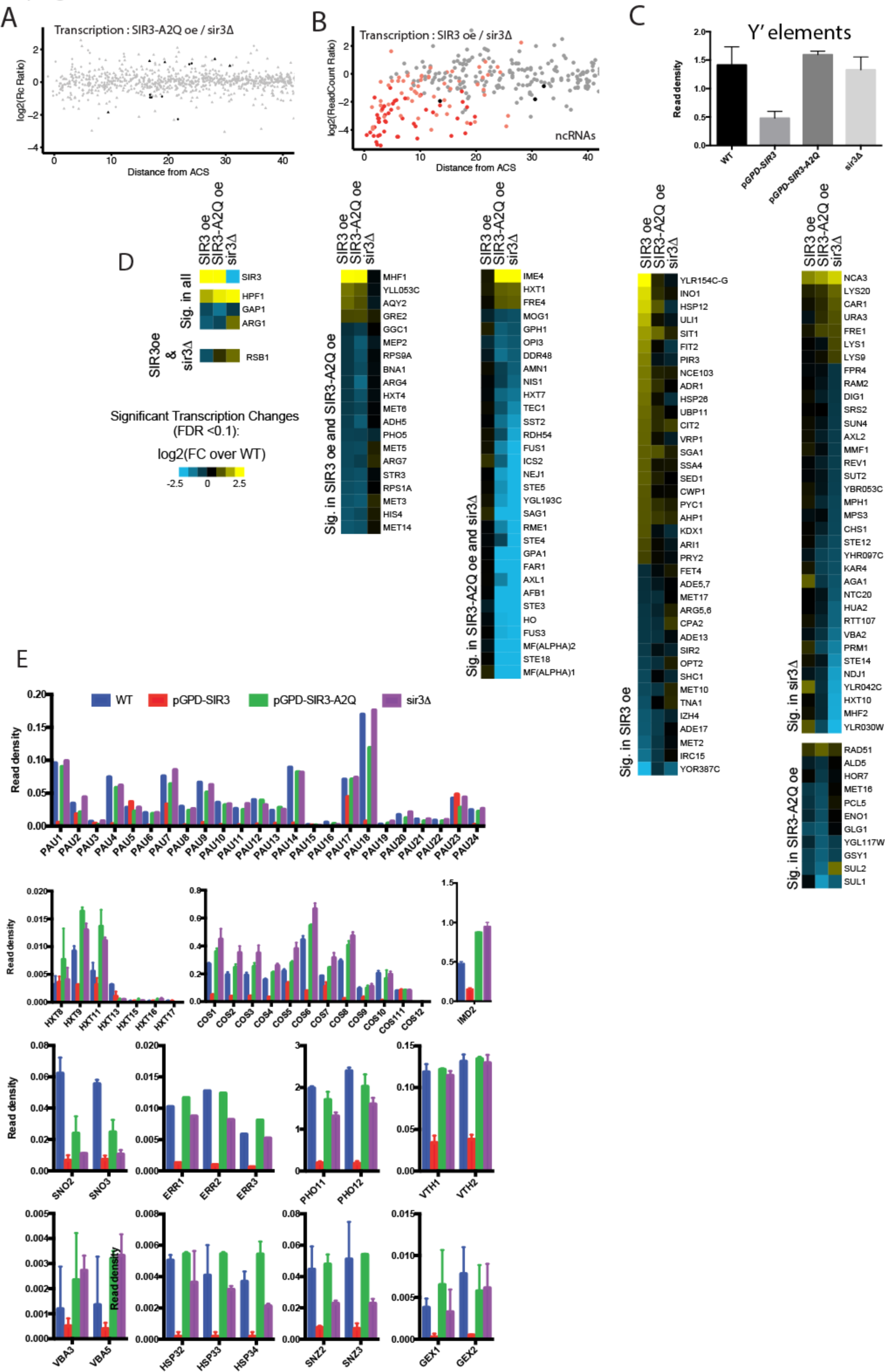
(A) Transcriptional changes in *sir3-A2Q* mutants versus *sir3* mutants. (B) Transcription of ncRNAs within subtelomeres, color code is identical to the main Fig2D. (C) Average Read density at Y' elements. (D) All transcriptional changes coined significant by EdgeR within euchromatin, color code indicates log2(FC). (E) Transcriptional changes of genes from subtelomeric families.

**Supplementary Figure 3:**
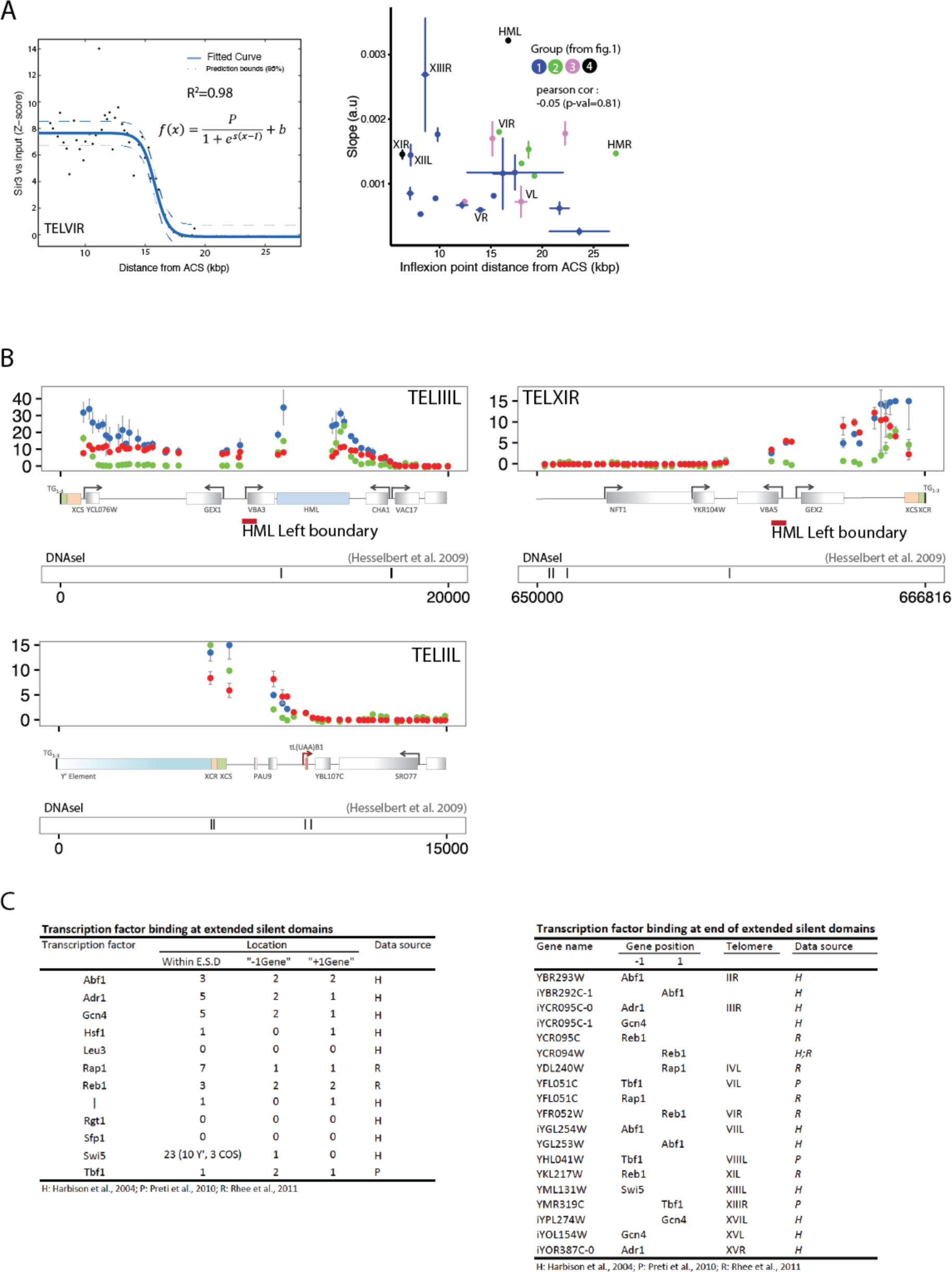
(A) Example of fitting of the ChIP-chip data, function used is shown on the graph. Right: Inferred slope versus position of inflexion point. (B) Examples of identified barrier at three subtelomeres at which Sir3 spreading does not extent when Sir3 dosage is increased. (C) Table listing transcription factor bound within E.S.D or at genes neighboring E.S.D. Original data source is indicated.

**Supplementary Figure 4:**
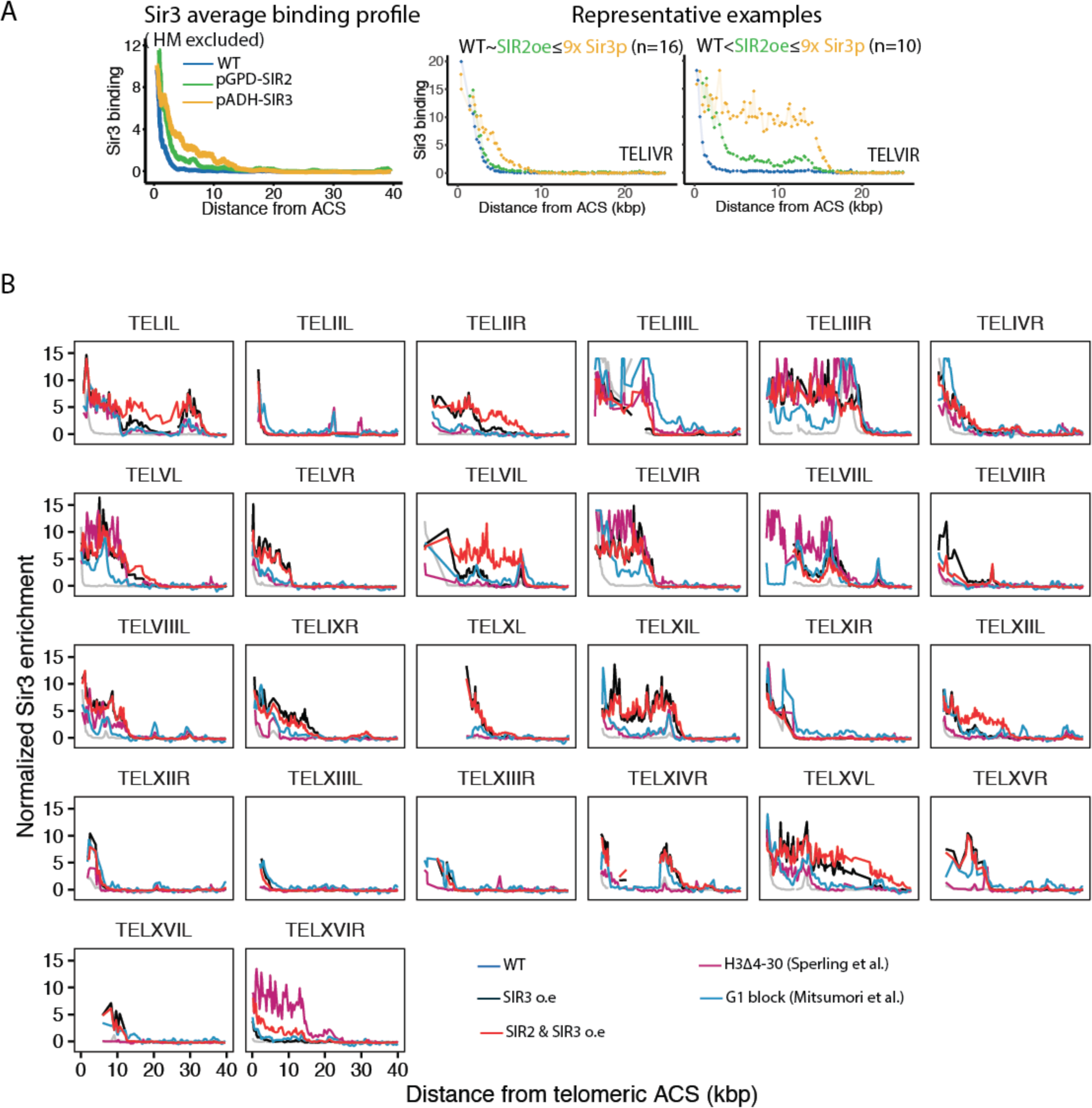
(A) Moving average of Sir3 binding at telomeres (with the exception of TELIIIL and TELIIIR, which contain HM loci) as in fig1F, in the indicated genotypes. Representative examples of Sir3 binding. Qualitative comparison of Sir3 spreading between conditions is indicated as legend with the number of subtelomere attributed to this stereotypical category. (B) Sir3 binding at individual subtelomeres. Enrichment corresponds to standardized Sir3 binding (z-score). Origin of external dataset is indicated. 6 Subtelomeres are not shown due to insufficient data.

**Supplementary Figure 5:**
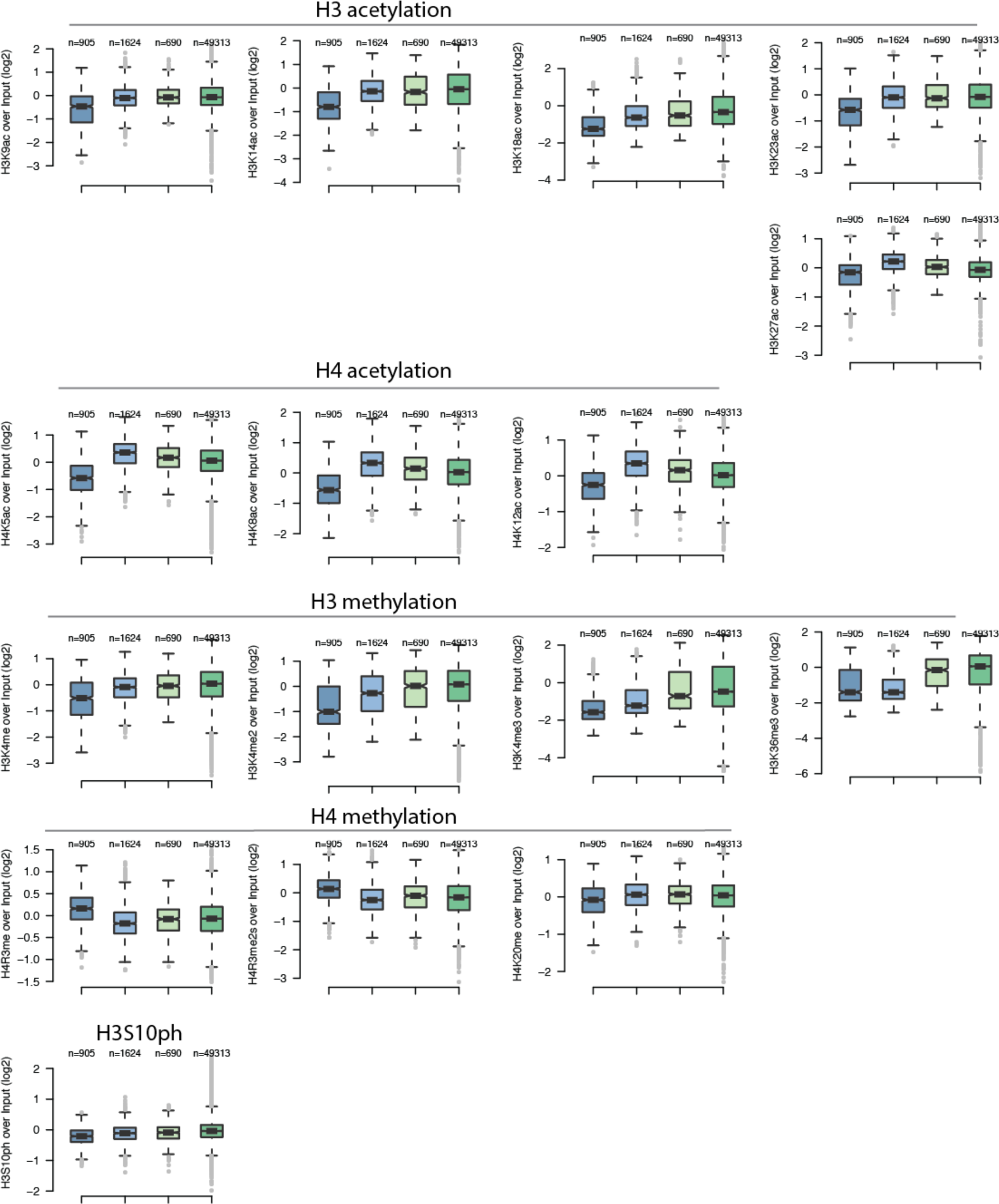
Distribution of selected histone marks relative to H3 (data from Weiner et al.2015) along wild type silenced domains and within the contiguous subtelomeric domains accessible to Sir3 upon overexpression. As a control, the distribution of those marks within the 5 kb contiguous to the end of extended silent domains as well as the genome wide distribution of those marks is shown.

**Supplementary Figure 6:**
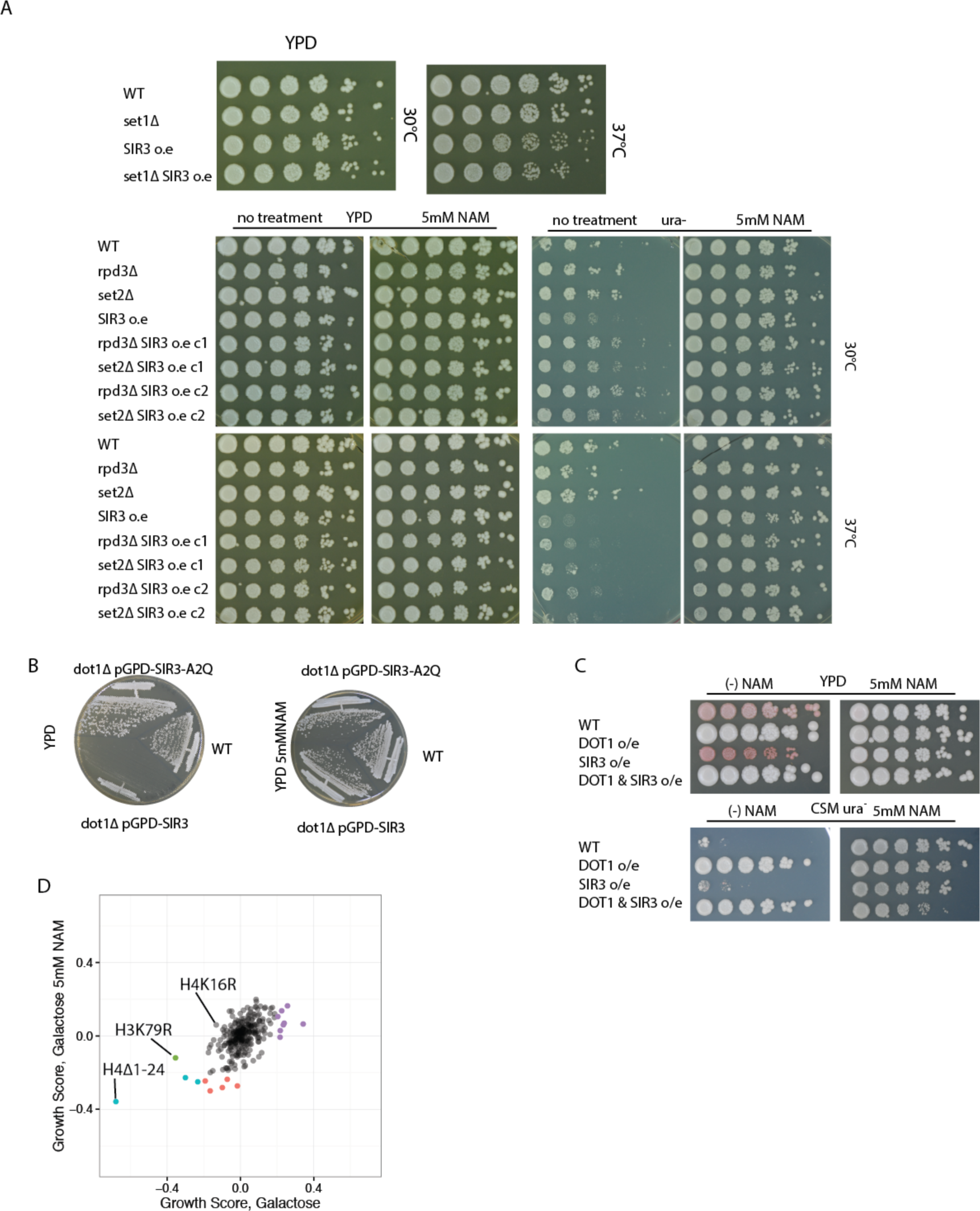
(A) Drop assays probing viability in the presence or absence of 5mM NAM. Protocol is identical to the one shown on the main figure. (B) *dot1* mutants overexpressing Sir3-A2Q are viable. (C) *DOT1* overexpression counteracts *SIR3* overexpression. WT strains have an *ADE2* reporter gene located at telomere VL. (D) SGA score of all histone point mutants probed, colored points pass our significance criterion and are colored according to their respective behavior (rescued by NAM treatment, sick in all conditions…).

**Supplementary Figure 7:**
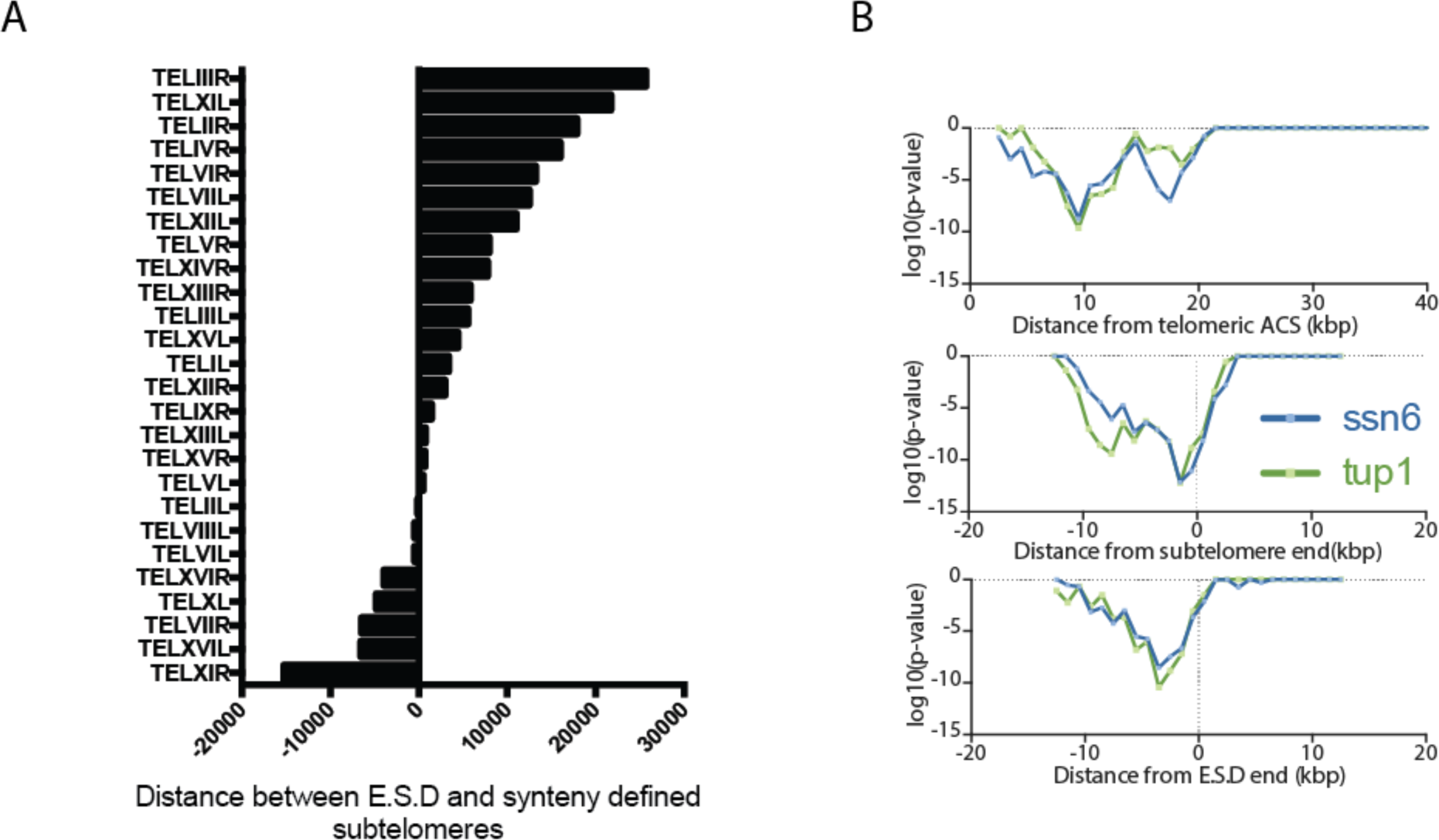
(A) Comparison of ends of E.S.D with subtelomere ends as defined by synteny in Yue et al. 2016. (B) Corrected p-values of hyper-geometric test for sliding 5kb windows (step=1kb) is shown for *tup1* and *ssn6* mutants in function of different subtelomeric viewpoints. This analysis corresponds to main figure 7B. Each point represents the center of a 5kb window.

**Strain table 1.**

Strain table
| ID | matingT. | genotype | background |
| --- | --- | --- | --- |
| 191 | alpha | ade2-1::ADE2 adh4::URA3-4xUASG-(C1-3A)n ppr1Δ::HIS3<br>rap1::GFP-RAP1(LEU2) | (W303) |
| 1254 | alpha | ade2-1::ADE2 adh4::URA3-4xUASG-(C1-3A)n ppr1Δ::HIS3<br>rap1::GFP-RAP1(LEU2) sir3::GPD-SIR3(NAT) | (W303) |
| 1256 | alpha | ade2-1::ADE2 adh4::URA3-4xUASG-(C1-3A)n ppr1Δ::HIS3<br>rap1::GFP-RAP1(LEU2) sir3::GPD-sir3-A2Q(NAT) | (W303) |
| 2487 | a | ade2-1::ADE2 hmlΔ::HPH rap1::RAP1-GFP(LEU2) | (W303) |
| 2627 | a | ade2-1::ADE2 hmlΔ::HPH rap1::RAP1-GFP(LEU2) sir3::pADH-SIR3(NAT) | (W303) |
| 2629 | a | ade2-1::ADE2 hmlΔ::HPH rap1::RAP1-GFP(LEU2) sir3::pTEF-SIR3(NAT) | (W303) |
| 2554 | a | ade2-1::ADE2 hmlΔ::HPH rap1::RAP1-GFP(LEU2) sir3::GPD-Sir3(NAT) | (W303) |
| 1667 | a | RAD5+ rap1::RAP1-GFP(LEU2) RDN1::ADE2 sir2::GPD-SIR2(KanMX) | (W303) |
| 1668 | a | RAD5+ rap1::RAP1-GFP(LEU2) RDN1::ADE2 sir2::GPD-<br>SIR2(KanMX)sir3::GPD-SIR3(NAT) | (W303) |
| 779 | a | ade2-1::ADE2 sir3::SIR3-GFP(LEU2) | (W303) |
| 3441 | a | ade2-1::ADE2 sir3::KAN pADH-SIR3-GFP(LEU2) | (W303) |
| 3442 | a | ade2-1::ADE2 sir3::KAN pTEF-SIR3-GFP(LEU2) | (W303) |
| 3443 | a | ade2-1::ADE2 sir3::KAN pGPD-SIR3-GFP(LEU2) | (W303) |
| 2056 | a | can1::MFA1pr-HIS3 hht1-hhf1::NatMX4 hht2-hhf2::[HHTS-HHFS]*-<br>URA3 where H4WT | BY4733 |
| 2986 | a | can1::MFA1pr-HIS3 hht1-hhf1::NatMX4 hht2-hhf2::[HHTS-HHFS]*-<br>URA3 where H3Δ4-30<br>Rap1-GFP(LEU2) | BY4733 |
| 2987 | a | can1::MFA1pr-HIS3 hht1-hhf1::NatMX4 hht2-hhf2::[HHTS-HHFS]*-<br>URA3 where H3Δ4-30 Rap1-GFP(LEU2) pGPD-SIR3(NAT) | BY4733 |
| 2476 | a | rap1::GFP-RAP1(LEU2) sir3::GPD-SIR3(NAT) | BY4741 |
| 3004 | alpha | ade2-1::ADE2 adh4::URA3-4xUASG-(C1-3A)n ppr1Δ::HIS3<br>rap1::GFP-RAP1(LEU2) bre1Δ::KanMx | (W303) |
| 3123 | alpha | ade2-1::ADE2 adh4::URA3-4xUASG-(C1-3A)n ppr1Δ::HIS3<br>rap1::GFP-RAP1(LEU2) bre1Δ::KanMx SIR3::pGPD-SIR3 (NatMx) | (W303) |
| 3180 | alpha | ade2-1::ADE2 adh4::URA3-4xUASG-(C1-3A)n ppr1Δ::HIS3<br>rap1::GFP-RAP1(LEU2) dot1Δ::KanMx | (W303) |
| 3181 | alpha | ade2-1::ADE2 adh4::URA3-4xUASG-(C1-3A)n ppr1Δ::HIS3<br>rap1::GFP-RAP1(LEU2) set1Δ::KanMx | (W303) |
| 3182 | alpha | ade2-1::ADE2 adh4::URA3-4xUASG-(C1-3A)n ppr1Δ::HIS3<br>rap1::GFP-RAP1(LEU2) dot1Δ::KanMx pGPD-SIR3-A2Q (NAT) | (W303) |
| 3183 | alpha | ade2-1::ADE2 adh4::URA3-4xUASG-(C1-3A)n ppr1Δ::HIS3<br>rap1::GFP-RAP1(LEU2) dot1Δ::KanMx pGPD-SIR3 (NAT) | (W303) |
| 3184 | alpha | ade2-1::ADE2 adh4::URA3-4xUASG-(C1-3A)n ppr1Δ::HIS3<br>rap1::GFP-RAP1(LEU2) set1Δ::KanMx pGPD-SIR3(NAT) | (W303) |
| 2838 | alpha | ade2-1::ADE2 adh4::URA3-4xUASG-(C1-3A)n ppr1Δ::HIS3<br>rap1::GFP-RAP1(LEU2) rpd3Δ::KanMx | (W303) |
| 2841 | alpha | ade2-1::ADE2 adh4::URA3-4xUASG-(C1-3A)n ppr1Δ::HIS3<br>rap1::GFP-RAP1(LEU2) rpd3Δ::KanMx pGPD-SIR3(NAT) | (W303) |
| 3301 | 1N | ade2-1::ADE2 adh4::URA3-4xUASG-(C1-3A)n ppr1Δ::HIS3<br>rap1::GFP-RAP1(LEU2) set2Δ::KanMx | (W303) |
| 3333 | 1N | ade2-1::ADE2 adh4::URA3-4xUASG-(C1-3A)n ppr1Δ::HIS3<br>rap1::GFP-RAP1(LEU2) set2Δ::KanMx sir3::GPD-SIR3(NAT) | (W303) |

